# Enabling AI in Synthetic Biology through Construction File Specification

**DOI:** 10.1101/2023.06.28.546630

**Authors:** Nassim Ataii, Sanjyot Bakshi, Yisheng Chen, Michael Fernandez, Zihang Shao, Zachary Scheftel, Connor Tou, Mia Vega, Yuting Wang, Hanxiao Zhang, Zexuan Zhao, J. Christopher Anderson

**Affiliations:** Department of Bioengineering, University of California, Berkeley, CA 94720, USA; QB3: California Institute for Quantitative Biological Research, University of California, Berkeley, CA 94720, USA; Physical Biosciences Division, Lawrence Berkeley National Laboratory, Berkeley, CA 94720, USA

## Abstract

The Construction File (CF) specification establishes a standardized interface for molecular biology operations, laying a foundation for automation and enhanced efficiency in experiment design. It is implemented across three distinct software projects: PyDNA_CF_Simulator, a Python project featuring a ChatGPT plugin for interactive parsing and simulating experiments; ConstructionFileSimulator, a field-tested Java project that showcases ‘Experiment’ objects expressed as flat files; and C6-Tools, a JavaScript project integrated with Google Sheets via Apps Script, providing a user-friendly interface for authoring and simulation of CF. The CF specification not only standardizes and modularizes molecular biology operations but also promotes collaboration, automation, and reuse, significantly reducing potential errors. The potential integration of CF with artificial intelligence, particularly GPT-4, suggests innovative automation strategies for synthetic biology. While challenges such as token limits, data storage, and biosecurity remain, proposed solutions promise a way forward in harnessing AI for experiment design. This shift from human-driven design to AI-assisted workflows, steered by high-level objectives, charts a potential future path in synthetic biology, envisioning an environment where complexities are managed more effectively.

## Introduction

Construction File (CF) is a domain-specific representation that encapsulates a genetic engineering experiment in terms of molecular biology operations and the genetic materials involved. Rather than being a language, it serves as an abstraction that defines the minimal information content necessary to describe the DNA modification chemistry involved in fabricating a DNA or genetic library. Despite the existence of multiple ways to express an experiment as a CF, we have explored its standardization to enhance communication among humans, software tools, and intelligent systems within a collaborative workspace. We propose specifications for two representations of CF: a shorthand format for convenience and a JSON version for cross-software communication. Furthermore, we provide parsers and simulators in Python, Java, and JavaScript. We explore human user interfaces for working with CF objects as well as AI interfaces and their ability to reason about CF objects.

Over the past years, numerous tools have been developed to assist with the design of DNA cloning schemes, such as J5^1^, Benchling^2^, A Plasmid Editor (ApE)^3^, SnapGene^4^, SBOL^5^, Biopython^6^, Geneious^7^, pydna^8^, GenoCAD^9^, and poly^10^. These tools have varying abilities to plan recombinant DNA experiments including the design of oligonucleotides and prediction of the resulting products. CF can serve as a standardized representation of the outcome of these design processes. It explicitly captures the experimental steps and their associated parameters in a minimal form independent of a specific software tool or environment.

The CF Shorthand Specification is much like a recipe for constructing DNA in the lab. A list of reaction steps is written in the order they should be performed, each defined by an operation keyword and parameters, separated by spaces. For instance, a Polymerase Chain Reaction could be specified as “PCR ForwardPrimer ReversePrimer Template ProductName”, with parameters representing names of DNA sequences or other relevant details. These sequences can be expressed in the CF as a name and sequence pair, like “T7_Universal TAATACGACTCACTATAGGG”, or they can reference DNAs from an external source such as a database. Although a CF does not specify implementation details such as the executor of the process (human or robot), reagent volumes, or manufacturer choices, it is still capable of defining the product sequences that would result from any successful implementation.

We first publicly introduced a format for CF in 2007 as part of a cloning tutorial on OpenWetWare^11^, with the intention of it being a human-readable representation of the experiment to aid in training and documentation. Over time, it became a practical necessity to develop software that could verify CF and catch design errors in these documents to avoid wasted lab resources and time. This need prompted multiple iterations of refining the ontology and syntax of CF, culminating in the current specification. Herein we provide multiple examples of CF shorthand that have been verified in the wetlab. We also present software tools that can read and simulate CF to ensure its completeness, syntactic correctness, and the feasibility of the proposed chemistry.

A CF can also function as an input or specification for an experiment, executable by an individual researcher, a core facility, or robotic systems. Although this paper does not present software for converting a CF into more detailed plans, it demonstrates that artificial intelligence can expand such a plan for human implementation. However, the current AI falls short of translating a CF into an Autoprotocol^12^, a JSON-based language that describes experimental procedures in terms of robotic operations, such as liquid transfers, plate sealing and unsealing, among others. Despite these limitations, there is potential for developing software that can perform this translation. Therefore, a CF can serve as a pivotal intermediate representation in the design process, with the remaining details inherently predetermined, provided a rubric that defines the resources available in the lab where it will be executed. This underscores the role of the CF as a critical intermediary in enabling intelligent systems, including AI, to effectively participate in the genetic engineering process.

## Results

The Construction File (CF) provides a structured framework for encoding genetic engineering experiments. This framework is articulated through two distinct specifications: a JSON object format (cf_JSDoc_specification.md) for precise machine-readable communication, and a shorthand format (cf_shorthand_specification.md) for human-readable documentation and quick notation. These specifications enable the encoding and decoding of experiment design information and lay the groundwork for the integration of artificial intelligence in experiment planning and simulation.

The JSON object format is a detailed representation. It consists of two main elements: ‘steps’ and ‘sequences’. The ‘steps’ element is an array of objects representing construction steps, including the associated operation, input sequences, and output product. The ‘sequences’ element is an object with key-value pairs, where each key represents a unique identifier for a DNA sequence, and the corresponding value represents the sequence, strandedness, and end chemistry of the DNA.

The shorthand format, on the other hand, is a more abstract and flexible representation. It is defined as a list of Steps, where each Step represents a specific operation in a molecular biology experiment. Steps are written on separate lines, with parameters separated by whitespace (preferably TSV). A Step includes the names of input DNA sequence(s), non-sequence parameters, and concludes with the name of the product DNA sequence. The input sequences can refer to products from previous steps. The shorthand format also allows integration of comments and sequences using ‘name sequence’ lines. This flexibility enables CF Shorthand to represent various DNA operations beyond those explicitly defined in the specification. However, parsers and simulator algorithms typically require a defined scope of operations and parameters to apply domain logic. To address this, level 1 of the specification specifically defines PCR, GoldenGate, Gibson, Digest, Ligate, and Transform operations.

As shown in Figure 1, the CF Shorthand provides a structured, machine-readable alternative to traditional illustrations of cloning strategies. Each step in the Construction File Shorthand begins with an operation, followed by operation-specific inputs, often sequence names. The final token in each step denotes the product, encapsulating the outcome of the operation. The full text of this CF is also available as Examples/Construction_pSB1A2-Bca9128.txt.

**Figure 1.**
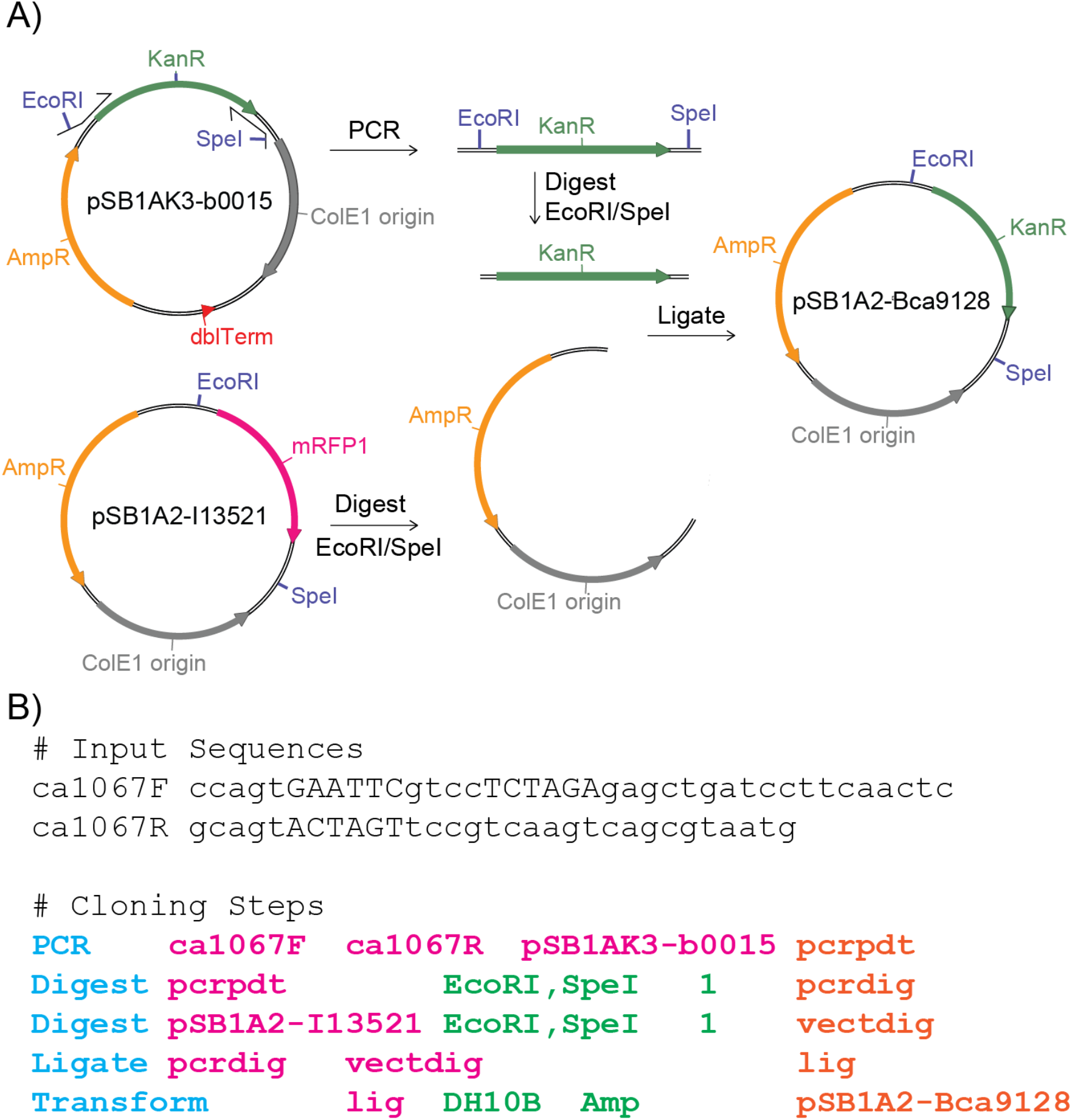
Shorthand Representations of a Cloning Strategy. (A) Conventional illustration of a cloning strategy, visually detailing PCR, Digestion, and Ligation steps. (B) Equivalent strategy represented in Construction File Shorthand. Each step begins with an operation (blue), followed by operation-specific inputs, often sequence names (magenta). The final token in each step (orange) denotes the product, encapsulating the outcome of the operation. This shorthand format provides a structured, machine-readable alternative to traditional illustrations.

Although the CF Shorthand format and the JSON format have different syntax, the only functional difference between the two formats is the level of detail regarding strandedness and other characteristics of the DNAs. In most real-world scenarios, cloning experiment inputs are either double-stranded DNAs longer than 100 bp or single-stranded linear oligonucleotides shorter than 100 bp. Consequently, the additional fields needed to express a DNA’s full structure can usually be inferred. One advantage of the shorthand format is its bidirectional compatibility with spreadsheets. Excel and Google Sheets can handle TSV data, allowing for easy manipulation and maintaining the TSV syntax when transferred between a text field and spreadsheet cells.

### Considerations for the specification

The CF specification was designed with a balance between detail and simplicity in mind. One approach could have been to describe steps in terms of lists of reagents, aligning with wetlab automation ontologies. However, this would have led to an unnecessary over-specification and would have been more difficult to simulate due to the need for a mechanistic simulation of each enzymatic step. On the other hand, a more abstract approach, aligning with standard assembly schemes like BioBricks and MoClo are simple to simulate, but this approach lacked the required detail for comprehensive representation of the diversity of experiments that are frequently performed. We also considered abstractly defining PCR to include mechanistically similar methods like Polymerase Chain Assembly and SOEing. However, this resulted in a heterogenous input parameter schema, leading us to define the operations more narrowly. A similar thing happened with Assembly. We explored an ‘assemble’ operation, and Gibson was an option for the enzyme. This abstraction didn’t add anything, and having assembly methods explicitly stated as operations was more direct. Thus, we selected commonly-used, method-level abstractions, encompassing the operations PCR, Digest, GoldenGate, Ligate, Gibson, and Transform. Each of these operations, in turn, have their unique requirements and parameters.

Beyond the naming of these operations, some require specific non-DNA parameters. For instance, the PCR operation includes an optional product size parameter, which is important when using the CF as an input specification. However, it is defined as optional since the PCR product size is unknowable if the PCR hasn’t already been simulated. Similarly, the Digest operation includes a ‘fragSelect’ index parameter. This specifies the fragment desired after digestion, with numbering starting from the first cut of the first enzyme. This approach offers flexibility and simplicity, as in most cases, the desired fragment is number 1. Finally, the Transform operation has an optional incubation temperature field that should only be included when it is a relevant detail. To further enhance flexibility and portability, sequences in the CF are treated in a specific way.

In the CF, sequences are referenced by their names, not as objects. This loose coupling allows a CF to be syntactically valid before the sequences associated with the names have been defined, thus allowing a CF to also serve as a specification for the design of the sequences. It also allows a CF to have alternate input sequences injected during simulation such that a similar sequence of cloning steps can be applied to different input DNAs. Additionally, it improves portability since memory-intensive sequence data does not need to be transferred.

In developing the sequence representation for the CF, we considered several formats including TSV, FASTA, Dseqrecord^9^, and a custom class, Polynucleotide. The simplest option, name and sequence of the ‘watson’ strand, was adopted for the shorthand format. For the JSON representation, we opted for a more detailed Polynucleotide object, capturing sticky ends, 5’ modifications, strandedness, and circularity. This representation, as illustrated in Figure 2, reflects the DNA’s state as it undergoes operation-specific transformations to yield expected products. This format accommodates atypical DNA forms and aligns with how molecular biologists often describe sticky ends. We also considered a Dseqrecord-like format wherein both strands of the DNA are expressed as strings along with an overhang integer. This offers chemical precision but requires additional processing and complex operations for AI reasoning. Moreover, the pydna implementation of Dseqrecord, while comprehensive, carries unnecessary complexity for our purposes and does not express 5’ modification chemistry. It also includes many fields inherited from Biopython’s SeqRecord about semantics and annotations which are not needed to specify the chemistry. A middle-ground representation, specifying whether the DNA is a plasmid, a dsDNA, or an oligo, was also included in shorthand. This covers most real-world scenarios and can be readily compiled to the Polynucleotide form.

**Figure 2.**
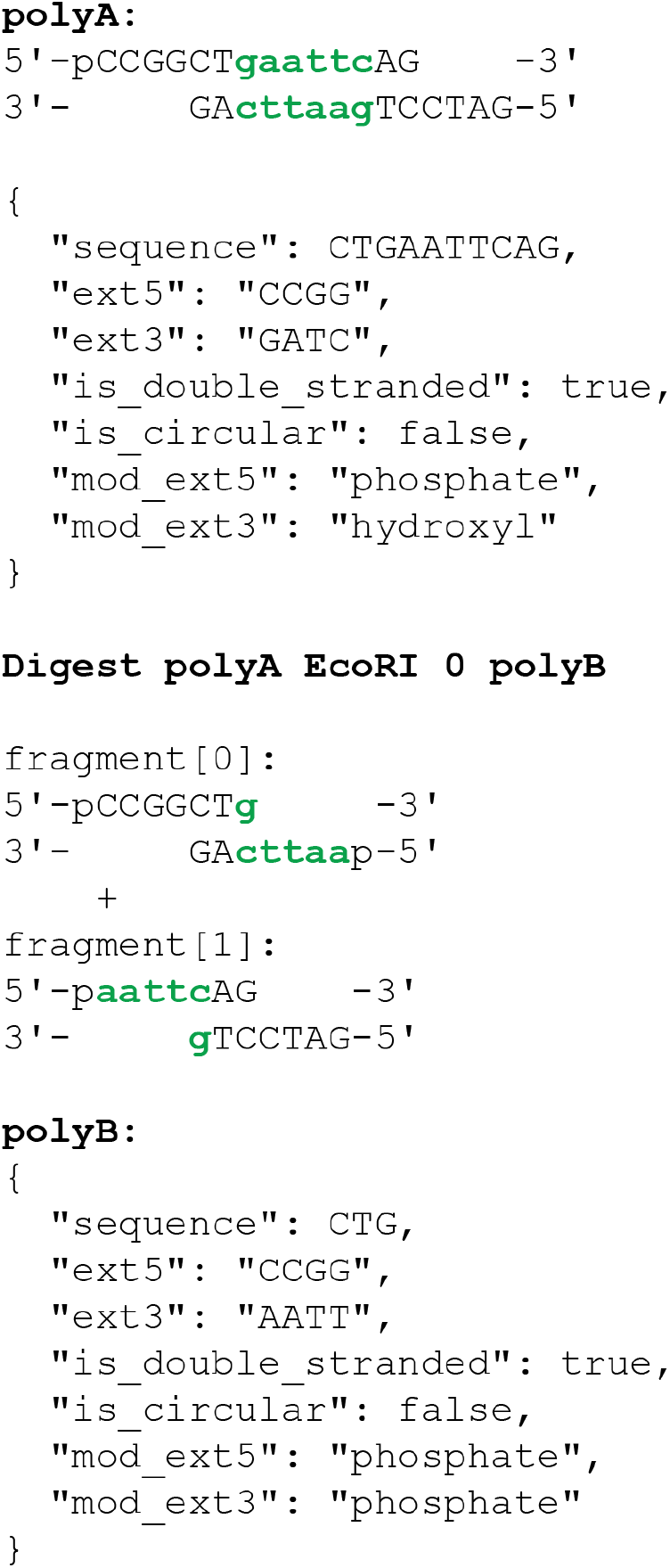
Polynucleotide Object Representation for Simulating Molecular Biology Operations. The hypothetical DNA ‘polyA’ is a linear, double-stranded DNA previously cut with BamHI, dephosphorylated, and subsequently cut with XmaI. In the Polynucleotide object representation, the fully duplexed DNA portion is captured as the “sequence”. Single-stranded overhangs are represented by the coding strand sequence as ext5 and ext3, denoting the overhangs on the left and right of the diagram, respectively. Modifications at the ends are indicated by enumerated types as mod_ext5 and mod_ext3. The simulation of an EcoRI digestion of this DNA would yield two fragments, indexed as 0 and 1. The ‘fragmentSelection’ field of the shorthand statement is set to 0, resulting in ‘polyB’ being returned as depicted. In the simulation software, Polynucleotides serve as dynamic representations of DNAs, reflecting their states as they undergo operation-specific transformations to yield expected products. Simulation software currently supports PCR, Digest, Ligate, GoldenGate, Gibson, and Transform operations.

**Figure 3.**
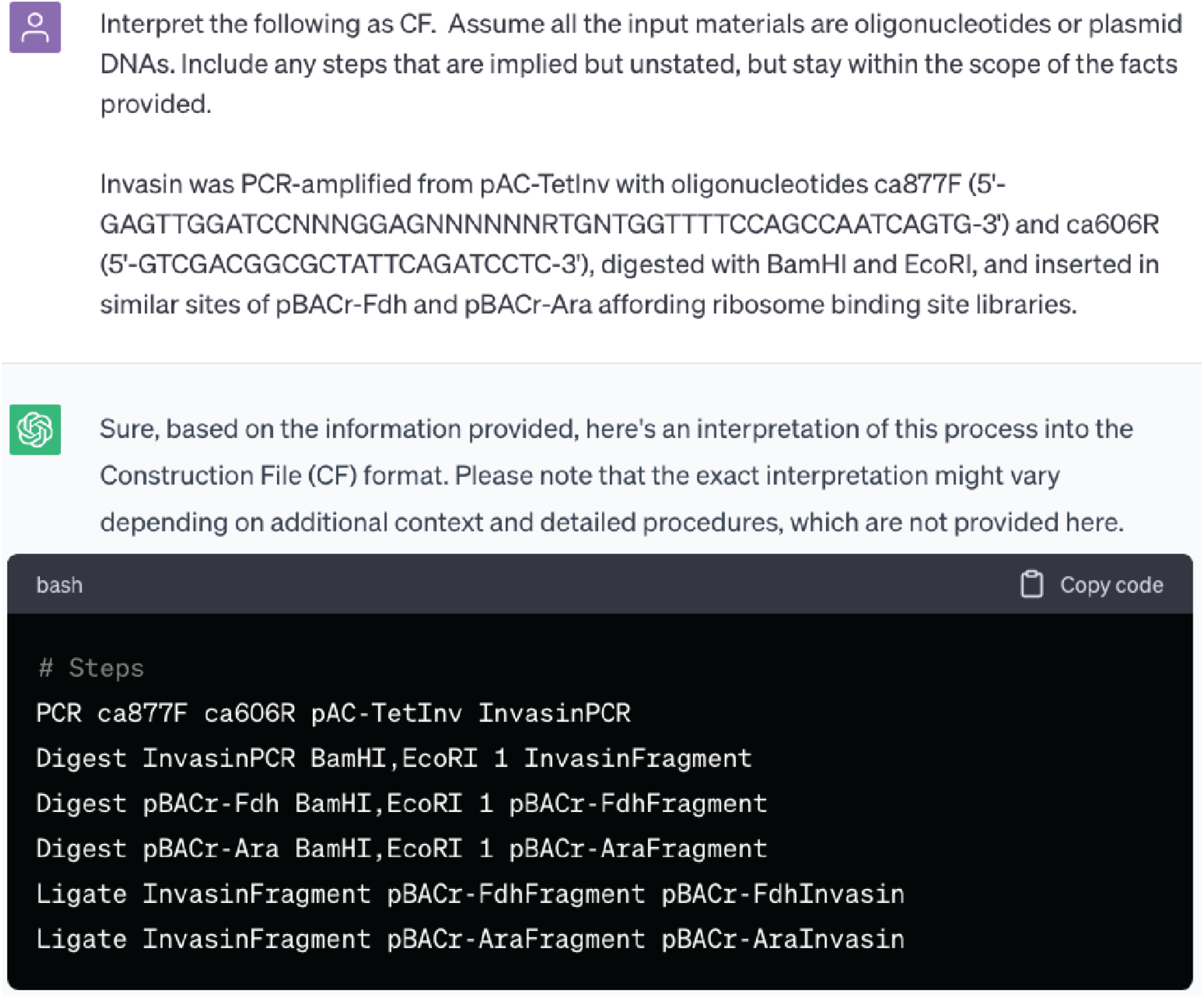
Zero-shot Natural Language Processing Interpretation of Construction Files by ChatGPT. After being prompted with the shorthand specification document, ChatGPT (GPT-4) demonstrates its ability to interpret plasmid construction text from a scientific paper into a construction file with high accuracy. This demonstration underscores the potential of A.I. to automatically extract construction files from scientific literature, opening new possibilities for large-scale, automated analysis of genetic engineering experiments from unstructured archival text.

### Assessment of AI in Interpreting, Designing, and Simulating CF

We conducted a series of experiments to assess the capabilities of AI, specifically GPT-4 via ChatGPT^13^, in interpreting, converting, and simulating CF. These experiments serve as an initial exploration of how AI can be integrated into the process of designing genetic engineering experiments. In each experiment, the shorthand specification text was provided at the start of the chat. The full transcripts of these chats are available as supplemental information under ‘Chats’, or via URL.

ChatGPT demonstrated a remarkable ability to interpret complex scientific text and convert it into CF shorthand. For example, when presented with a published description of a cloning experiment involving the preparation of two ribosome binding site libraries^14^, ChatGPT accurately interpreted the steps and converted them into CF shorthand, despite the complexity of the experiment and the need to infer unstated steps from the text (invasin_parse_test.html). This result suggests that a literature mining effort to extract the history of published recombinant experiments is within reach of current technology, although it is beyond the scope of this study.

We also explored if ChatGPT could perform zero-shot design of a CF. After providing the shorthand specification, we tasked it with performing a ‘prefix insertion’ on two BioBrick plasmids (design_biobrick.html). ChatGPT returned a syntactically correct CF, correctly inferring the need for two digestion reactions and one ligation reaction. However, it initially chose incorrect enzymes for the digests. After providing additional information from an external website, ChatGPT corrected the enzymes and structure in the CF. The only remaining error was the ambiguity of the fragmentSelection indices, which was resolved with further prompting about the orientation of the input sequences. This experiment demonstrated that, with corrective prompting, GPT can be guided to author accurate construction files.

Interconversion between different forms of CF is another area where ChatGPT showed proficiency (syntax_conversions.html). Given the specifications for shorthand and JSON formats, it was able to convert a CF from shorthand to JSON, correctly inferring the strandedness and circularity details for the DNAs involved (Examples/Construction_pSB1A2-Bca9128.json). We also asked it to generate an XML version (Examples/Construction_pSB1A2-Bca9128.xml), demonstrating the flexibility of CF and the ability of GPT to handle different formats.

The generation of human-readable work plans and Autoprotocols from CFs is a more complex task, and here ChatGPT showed both its capabilities and limitations. When asked to reduce a CF to a work plan that could be passed to a technician (technician_instructions.html), ChatGPT produced mostly correct instructions. However, it hallucinated locations for preexisting samples and omitted some steps that are typically included in such instructions, such as full calculation of the reagent volumes and consideration of DNA concentrations. When asked to generate an Autoprotocol, a JSON-based language for robotic liquid handlers, ChatGPT struggled (autoprotocol_instructions.html). Despite being familiar with Autoprotocol, it was unable to produce valid JSON, indicating that the leap from CF to Autoprotocol is currently beyond GPT’s capabilities.

Simulating CFs directly in ChatGPT also presented challenges. When given the entire text of a CF, the token limit was exceeded due to the long length of plasmid sequences. Shortening the sequences in the CF allowed ChatGPT to accept the prompt, but it failed to simulate the result due to the task’s complexity (invasin_simulation.html). Thus, while GPT shows promise in understanding and interconverting CF, it struggles to accurately design, simulate, or compile them into wetlab instructions. Given the paramount importance of accuracy for BioCAD tools, these findings underscore the need for a more precise approach, such as could be achieved with a GPT plugin.

### PyDNA_CF_Simulator: a Python-Based ChatGPT Plugin for CF Simulation Using PyDNA

To explore the possibility of GPT directly invoking python scripts for simulation tasks, we attempted to have GPT generate a pydna script representing the pSB1A2-Bca9128 example CF (cf_to_pydna.html). The pydna library shares a similar ontology with CF and includes simulators for PCR, digestion, ligation, and Gibson assembly methods. However, the resulting script from GPT required us to make several manual adjustments, including moving the pip statement, adding the DNA sequences, and removing the API requests to GenBank. Despite these corrections, GPT incorrectly used an ‘Assembly’ function to simulate ligations rather than the ‘+’ operand on the sequence objects, rendering the script unrunnable. This experiment led us to conclude that GPT’s current capabilities are insufficient for writing this executable representation of CF.

While Python scripts are useful, they present several challenges when used as documentation for construction files in an AI interface. Firstly, they are written in free-form Python, there are potential security issues with an interface that executes these scripts. Secondly, they assume a specific software implementation, limiting extensibility and interoperability in a multi-tool environment. Lastly, Python scripts do not readily enable inspection, a crucial feature for using CF as a specification.

To address these limitations, we developed a Python plugin wrapper, PyDNA_CF_Simulator^15^, capable of parsing CF and executing the appropriate pydna syntax for simulation. We created Python classes for ConstructionFile and Polynucleotide according to the jsDoc spec, and developed functions for parsing Strings of CF shorthand or JSON into these classes. Functions were also created to interconvert between Polynucleotide and Dseqrecord representations. We then developed a function that simulates a ConstructionFile instance, executing the appropriate operations and returning the resulting product sequences. Finally, we created an API wrapper to host the simulator as a REST endpoint, along with a YAML and manifest containing the shorthand specification for communication with ChatGPT.

Testing of the Python plugin wrapper revealed several limitations. While the plugin successfully handles simple cases like PCR on short templates (pydna_plugin_test.html and pydna_plugin_test.mov), its token limit in the low thousands significantly curtails its utility with larger DNA sequences. This limit is far from sufficient to encode complex structures like plasmid sequences, let alone the millions+ tokens required to express a genome sequence. Due to its limited utility, we have not submitted PyDNA_CF_Simulator for inclusion as an official ChatGPT plugin. However, the code is available on Github under the open-source MIT license.

Further limitations were found within the pydna library itself. Pydna’s inability to simulate Golden Gate reactions, a cornerstone of modern synthetic biology, greatly restricts its utility for a wide range of experiments. Although the source code includes a script for it, it is not fully implemented. While Golden Gate could be described as sequential digestion and ligation steps, which are implemented, this is not equivalent to the simultaneous cutting and ligation that occurs in the actual process which requires additional logic. Additionally, pydna allows non-DNA letters, even permitting the entire alphabet as syntax.

While the Python plugin wrapper effectively delegates the simulation task to reliable, well-tested code, it has notable limitations. A significant challenge with this type of interface is the absence of visualization and persistence for both the resulting sequences and the Construction File itself. Ideally, an additional interface would be integrated into the workflow to provide users with a clearer understanding of the process and its outcomes. These findings highlight the necessity for further development and enhancements to the AI interface, particularly in the areas of user interface design and strategies to circumvent token limits.

### ConstructionFileSimulator: a Java-Based Tool for Validation and Simulation of CF

There are two distinct types of software that could be developed for simulating Construction Files (CFs): one that validates the CF, and another that calculates the product. While these objectives may seem similar, they lead to different design decisions and implementations. For instance, consider a Golden Gate assembly of three fragments, where one fragment has compatible ends on both sides and thus will re-ligate. A tool focused on calculating the product would correctly simulate this scenario and return the single-fragment product. However, a tool focused on validating a CF would instead identify this scenario as a problem, alerting the user to the potential issue rather than simply returning the result. This focus on error detection and prevention is crucial for ensuring the validity and success of genetic engineering experiments.

With this validation objective in mind, we developed the first iteration of ConstructionFileSimulator (CFS) in Java^16^. We employed a programming style reminiscent of Functional Programming with mostly-pure functions and immutable classes. It interprets CF shorthand text into a ConstructionFile object, subsequently simulating the expected reaction product step by step. If an error arises during simulation, it triggers an error response which terminates the operation and delivers a detailed message to guide corrective action.

The relationship between CF operations and simulator functions in CFS is largely one-to-one, but the concurrent development of the CF syntax, CFS, and wetlab usage has led to the need for backward compatibility with past versions of CF. As a result, CFS can handle a broader array of syntax than the specified shorthand, and the codebase contains more complexity than strictly necessary. It also supports PCA (Polymerase Chain Assembly), SOE (Splicing by Overlap Extension), and Klenow (Klenow extension) operations which are not in the specification. From a system architecture perspective, it’s worth noting that a strict one-to-one correspondence between operations and functions is not always the most efficient or effective design. For example, lower-level functions such as reverse complementation (RevComp.java) are used across multiple algorithms and are therefore implemented as standalone functions rather than being associated with specific operations. Furthermore, to accommodate a variety of PCR-like scenarios, we generalized these techniques in the simulation. While this abstraction was challenging to express in shorthand, it provides a compact solution at the functional level. The CFS codebase also includes several exploratory and vestigial features that we have omitted from this discussion for the sake of focus.

Within this architecture, the project houses two PCR simulators, each designed to address specific experimental scenarios. The simpler one, encoded in the method perfect18Simulation, is only activated when a singular template and two oligos are present, with both oligos perfectly matching the template over 18 bp at their 3′ ends. This condition is usually met for standard cloning experiments. However, for non-standard scenarios, such as site-directed mutagenesis involving 20-mer oligonucleotides with a central mismatch, a more mechanistic simulation is needed. This includes simulating PCA, SOEing, or Klenow Extension, where template varieties from single-stranded to double-stranded, and their quantities from 0 to n, must be considered. To accommodate these scenarios, the PCRSimulator employs a backup algorithm, which mimics pairwise DNA annealing and extension. It checks for alignments where the 3′ six bases of the oligo exactly match the template, then uses JAligner and Tm calculations for further detection of annealing sites. However, this more complex function, while generally reliable, occasionally struggled with scenarios that a simpler algorithm could handle correctly. Additionally, it was computationally demanding, causing failures for longer templates and occasional inability to detect obvious annealing sites. To mitigate this, the simpler version is used as a first attempt before falling back to the more mechanistic simulation when necessary. The simulator can now handle more scenarios than outlined in the specification documents, including unique cases like mixtures of single-stranded and double-stranded templates. Both algorithms have been rigorously tested and confirmed to work on linear and circular templates, including inverse PCRs, and they handle 5′ modifications, 5′ extensions, and common issues such as multiple annealing sites and orientation errors.

The Digest operation uses a REBASE database-derived file for restriction enzyme information, making it capable of handling more enzymes than mentioned in the specification. It correctly handles degenerate cutters, both 5′ and 3′ extensions, and appropriately assigns phosphates to the 5′ modifications of freshly cut DNAs. Though there is a method in the code (cutOnce) that simulates a single cutting event, the Digest operation is assumed to mean ‘cut to completion’ and thus does not support partial digests.

In the simulation of ligation, the presence of a 5′ phosphate and matching sticky ends are checked, and two matching ends of two input Polynucleotides are concatenated into one. This process is repeated until only one fragment remains. If its ends are compatible, it is denoted as a circular DNA, and the sticky ends are integrated into the sequence field of the resulting Polynucleotide.

The simulation of GoldenGate primarily involves cutting with the type IIS enzyme and ligating the fragments, with additional checks for orientation, number of sites in the molecule, and the appropriateness of the generated sticky ends. Gibson simulation finds exact 20 bp matches between homologous ends and connects the DNAs pairwise.

The simulation of transformation, while implemented, is currently limited to checking that the product is circular. This is because transformation of a bacterium with a DNA requires it to be circular. However, it’s worth noting that CFS fully enables the generation of in vitro linear DNAs, which can be useful in certain scenarios, such as library fabrication.

CFS includes rigorous checks for possible design errors and provide comprehensive error messages when triggered. Over the course of three years, our use of the CFS for validating wetlab designs, along with its extensive application by over 100 students, has enabled us to identify and rectify numerous bugs. This iterative process led to the creation of a multitude of unit tests for various edge case scenarios, enhancing the reliability and robustness of our simulator. The development history is documented as issues in the ConstructionFileSimulator repository on Github.

CFS’s most straightforward interface is its SimulatorView Swing GUI, launched by executing the jar file without arguments. This interface accepts a construction file’s shorthand text and outputs the final step’s product. As illustrated in Figure 4, we fed the GUI with steps parsed by ChatGPT from the native invasin text, along with the sequences of the three input plasmid sequences (Chats/ invasin_cf.txt). The resulting sequence of pBACr-AraInvasin matches the expected map and aligns with sequenced isolates, validating the simulator’s accuracy and utility. We have provided an array of real-world examples (found in the supplemental Examples folder), showcasing the successful application of CFS. These examples feature experiments that involve degenerate bases, the creation of libraries, SOEing, PCA, and Klenow extension, all of which the simulator correctly handles.

**Figure 4.**
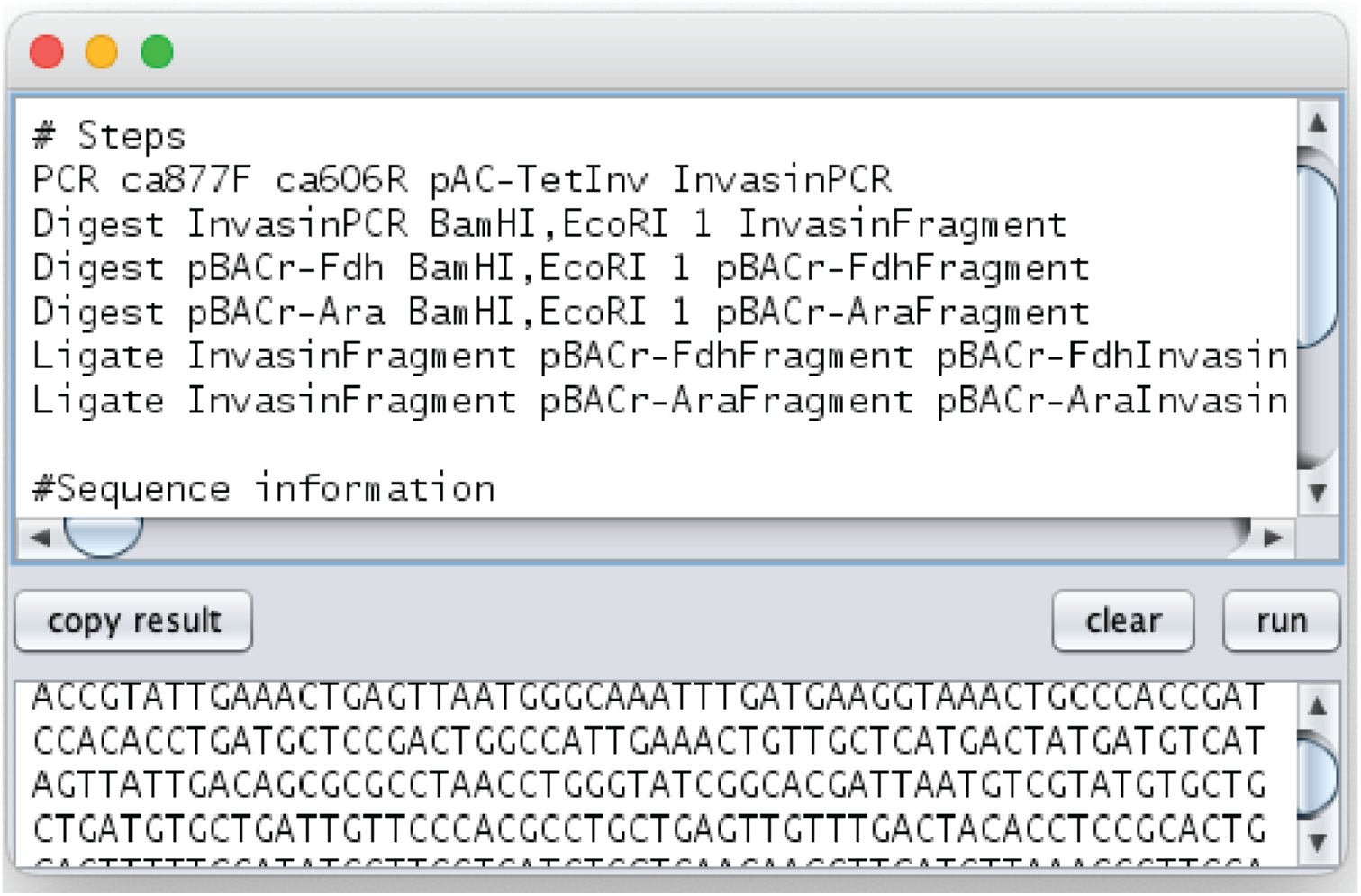
Simulation of Invasin Construction File in a script editor. SimulatorView, a simple GUI included with ConstructionFileSimulator, accepts the text of a construction file and outputs the product of the final step. In this instance, the GUI is provided with the steps parsed by ChatGPT, along with the sequences of the three input plasmid sequences. The complete document can be found in the supplementary file ‘invasin_cf.txt’. Upon clicking ‘run’, the construction file is simulated step-by-step. The resulting sequence of pBACr-AraInvasin aligns with the expected map and is consistent with sequenced isolates, demonstrating the accuracy and utility of the simulation.

### Mitigating Clerical Mistakes with ‘Experiment’ Objects in ConstructionFileSimulator

While simulating a CF is an effective way to detect technical errors in experimental design, such as oligo design issues, it doesn’t account for clerical errors that often occur in larger experiments involving multiple CFs or during collaborations among research teams. These errors, such as maintaining different versions of input sequences, are surprisingly common and can severely impact the success of an experiment. To address these issues, we’ve introduced the concept of an ‘Experiment’ object into CFS.

The creation of an ‘Experiment’ object begins by passing a hard drive path to a folder containing all relevant files to a parser. This includes sequence files in TSV or GenBank format (.gb, .seq, .str, and .ape), CFs expressed as plain text files, and additional sequence files, primarily for oligos, in a TSV format that also allows additional columns. This format is particularly useful as it aligns with the IDT oligo form, reducing the risk of error when copying and pasting between what is simulated and what is ordered. The parser then outputs an ‘Experiment’ object that encapsulates all the provided information. Once the ‘Experiment’ is created, it can be simulated to ensure the accuracy of the documentation.

Executing a list of CFs requires additional analysis to determine the correct order of execution. This is crucial to ensure that the products of earlier files can be used as inputs for later files. For example, in the pTP2_reporter example, a series of unrelated experiments was used to construct a reporter plasmid. To correctly simulate this, the software must identify the order of execution for each CF. We also had to consider the potential for name reuse, such as “pcrpdt” to refer to products of intermediate steps. To address this, the CFS implementation of ConstructionFile includes an explicit singular output from the entire file, which is set as the product of the last step during parsing. This addition, while not explicitly part of the specification, is necessary to resolve potential conflicts and implies that a ConstructionFile describes not only an ordered list of steps but also a specific product outcome.

The need for this higher-order ‘Experiment’ object is heavily dependent on the user interface. Our current approach treats files as contents of a folder, but other systems might use a database, potentially reducing the impact of clerical errors if the design and simulation functions were integrated. Furthermore, the exact content and format of an ‘Experiment’ are yet to be defined. Within the CFS, it encompasses sequences and CFs, but a more comprehensive specification could include measurement data, analysis, and more. Therefore, while this ‘Experiment’ functionality is part of the CFS project, we currently propose no standards for it and present it as an exploratory feature.

An ‘Experiment’ folder can be parsed and simulated using the SimulateExperimentDirectory function. This function is executed when the user runs the jar from the command line and passes in the path to the folder as a parameter. SimulatorView will also execute this function when such a folder is dragged-and-dropped on the GUI. Upon execution, the simulator generates a GenBank file for each product sequence and creates two log files: C5seqs.txt, which contains all sequences (inputs, intermediates, and products), and C5log.txt, which provides a detailed account of all events that occurred during execution. These log statements are also outputted to the command line when the jar is run from there. This information is helpful for identifying and correcting errors in the experiment’s design or documentation.

There are two supplemental examples of ‘Experiment’ folders that can be run with CFS. The Lycopene2 example demonstrates a scenario where several construction files are performed in parallel using a shared set of oligos in different combinations. The pTP2_reporter example illustrates a chain of sequential CFs where the product of one becomes an input to another. A demonstration of running CFS on this example is available at cfs_experiment.mov.

### C6-Tools: Simplifying CF Simulations and Oligo Design in Google Sheets

The Java implementation of CFS is well-tested, reliable, and effective for validating correct CF. However, students have found it somewhat challenging to identify errors in the CF. The log file details all events, which, while helpful in pinpointing errors, can result in a complex interaction akin to code debugging.

Typically, we run simulations through the IDE and leverage its debugging tools. A significant part of this challenge stems from the lack of visual representation during simulation. Although this issue could be addressed by creating more graphical user interfaces, this also presents another learning hurdle.

Driven by these usability concerns, we ventured into developing a variant of CFS, named C6-Tools^17^, using Google Apps Script (JavaScript) within a Google Sheet. In addition to simulation functions, we also integrated algorithms for oligo design. Displaying individual design and simulation events in a 2D spreadsheet grid significantly simplifies the visualization of ongoing operations and error identification. Additionally, this interface is highly familiar and requires little explanation for new users and can be easily accessed via url. While C6-Tools offers a lower entry barrier compared to CFS, it is a newer tool and has not been as extensively tested.

Initially, our aim in developing C6-Tools was to leverage GPT-4 to automatically translate the Java code into Apps Script. However, the majority of the functions proved this task to be not as straightforward. One notable complication was the Sheets’ inability to accept objects as cell values, necessitating their additional management as JSON. Despite these hurdles, ChatGPT was instrumental in facilitating this process with extensive prompting and revision.

Given the prevalence and versatility of Gibson Assembly and Golden Gate cloning in modern genetic engineering, traditional methods like digestion and ligation have become less relevant. Therefore, we decided not to include support for these older ‘cut and paste’ methods in C6-Tools. Additionally, this led to significant simplification of the code. Though Polynucleotide and its tracking of end chemistry is needed to simulate separate digestion and ligation steps, it is not needed to accurately simulate PCR, Golden Gate, and Gibson methods which can be well handled by simple sequence strings. Nevertheless, we include a class definition for Polynucleotide as an option for future development in JS.

Nonetheless, constraints such as the lack of a testing environment, the inability to import libraries, and the non-portability of the code hamper further development of C6-Tools in its current form. In the case of Apps Script, the scripts must be either within the Sheet file, requiring duplication, or transcluded from a library with a complete wrapper. This quality of Apps Script limits the ability to maintain and extend the library. However, for users who may wish to customize their own version of C6-Tools, the independence offered by the current implementation may be preferable over a development team’s oversight.

## Discussion

The CF specification, as it currently stands, covers a wide range of common molecular biology operations, including PCR, Digest, GoldenGate, Ligate, Gibson, and Transform. However, there are several common methods that are not explicitly represented in the CF specification. These include USER cloning, Ligase Chain Assembly (LCA), QuikChange mutagenesis, site-specific recombination systems like CRE/Lox and Gateway, homologous recombination methods like Datsenko-Wanner, and transposon-based methods. In addition, the CF specification does not currently support simple annealing of oligos to form a duplex DNA, CRISPR-mediated DNA cutting, or TOPO-TA cloning.

Each of these methods has unique requirements and parameters that would need to be incorporated into the CF specification to enable simulation. For example, QuikChange mutagenesis involves a PCR-like process, but the product is not the same as a typical PCR product. The CF PCR algorithm, while generalized, does not currently infer homology and reclosure of ends that QuikChange would require. Similarly, site-specific recombination systems like CRE/Lox and Gateway involve specific recognition sequences, which would need to be identified during the simulation process.

In addition to these specific methods, there are also broader categories of techniques that are not currently covered by the CF specification. For example, the CF specification does not currently support the representation of mixed pools of entirely different sequences, which are often used in library construction. Nor does it support the representation of more complex DNA structures, such as DNA bubbles or mixed RNA/DNA structures.

However, the question remains: do we need to include all these methods in the CF specification? The answer largely depends on the specific goals and use cases of the CF specification. If the goal were to create a comprehensive database of all cloning experiments, then a comprehensive representation of all possible methods would be necessary. On the other hand, if the goal is to provide a simple and intuitive interface for designing common molecular biology experiments, then a more limited set of operations may be sufficient. In any case, the decision to include or exclude specific methods from the CF specification should be made with careful consideration of the trade-offs between comprehensiveness, simplicity, and practical utility. Particularly for interacting with intelligent systems, where token limits are important constraints, having to include endpoints or API info about all these different operations gets heavy. If the user isn’t going to do all these things, then why are they in the tool?

### Broadening the Scope of the Transform Operation for Greater Experimental Accuracy

The Transform operation, a pivotal step in many molecular biology experiments, signifies a phase where the DNA is subject to further chemical modifications. For example, DNA nicks can undergo resealing, and the host’s dam and dcm systems can introduce novel methylation patterns. Nonetheless, the existing Transform operation neither accounts for these modifications nor verifies the presence of a selectable marker or a suitable origin of replication. Moreover, it does not confirm whether the replicon will replicate in the designated host, or if other plasmids originating from the same incompatibility group already exist in the strain. To fully validate a transformation, the Transform operation would need to incorporate these checks. Furthermore, the current ontology, with its *E. coli*-centric focus, presumes the use of antibiotics that may not be suitable for yeast work or transfection in plant or animal cells.

Beyond these fundamental verifications, the Transform operation could be enhanced to encompass a more exhaustive simulation of the biological processes initiated upon the entry of DNA into the cell. Such a simulator could scrutinize the introduced DNA for proper gene structure and assess the overall cellular system for biochemical accuracy, a first pass at which we presented in our previous work^18^. This would entail simulating the intracellular biochemical reactions and forecasting the cellular response to the introduced DNA. For example, the simulator could check if a sufficient grouping of genes was introduced to complete a pathway to a desired metabolite. It could infer promoter behavior and determine what regions of the DNA would be transcribed and translated. By juxtaposing this inferred pathway data with a functional specification, the simulator could ascertain whether the designed system would operate as intended, or if it could potentially be toxic to the cell or contain elements that might cross-react. Incorporating a transform simulator would provide an additional layer of validation, ensuring the precision of experimental designs and thereby enhancing the overall dependability and trust in the CF.

### Appraising User Interface Considerations for Effective CF Deployment

During the development of CF, we explored a range of interfaces, each presenting unique advantages and challenges. These interfaces include the Python scripting interface, the SimulatorView shorthand editor interface, the Experiment folder-based interface with CFS, the spreadsheet interface via C6-tools, the API interface with PyDNA_CF_Simulator, and the ChatGPT conversational interface.

The Python scripting interface provides a robust and flexible platform for designing and simulating experiments, a feature that programmers will find familiar. However, its accessibility is limited to those with coding experience. Conversely, the SimulatorView shorthand-script based interface is perfectly suited for crafting bespoke, detailed experiments. Yet, it may be cumbersome when handling many files due to its manual nature.

The spreadsheet interface, facilitated by C6-tools, offers the advantages of visual arrangement, lookup properties, and easy portability. It also enables the use of all the other spreadsheet functions, including the ability to drag the contents of a field across a range. This makes it particularly useful for describing sets of constructs, such as an ortholog or promoter scan.

The Experiment folder-based interface offers portability and compatibility with filesystem contexts like Github or Google Drive. It can be zipped and sent via email, making it a convenient option for sharing and collaborating on experiments. The API interface with PyDNA_CF_Simulator allows for programmatic interaction with the CF tools, providing another layer of flexibility.

It’s pertinent to highlight that the most common approach to authoring CF likely does not involve direct typing. Instead, a collection of design functions could generate the CF or Experiment object. Consider, for instance, an ortholog scan function. This function would take as input the initial prototype plasmid, specify the ORF to be scanned, and the organisms from which an ortholog is desired. The function would then execute a BLAST search of the ORF sequence to be replaced against the specified organisms, select an appropriate cloning strategy, design all necessary oligos, and output an Experiment object ready for simulation or execution. Preliminary versions of such algorithms are presented for oligo design in C6-tools, but we reserve the development of such functions for future work. Ultimately, a comprehensive library of such design functions could be established to cater to a wide array of scenarios.

The ChatGPT conversational interface facilitates a more intuitive interaction by leveraging natural language processing. However, it is currently limited by token limits, which restricts the complexity and length of the interactions. Ideally, the AI could be aware of all the other interfaces such that it could, for example, build a spreadsheet that invoked the functions, or translated a spreadsheet to the experiment folder format. This would allow the AI to leverage the benefits of each interface, while mitigating their individual limitations.

### Challenges and Opportunities in Integrating AI for Experimental Design

Synthetic biology stands on the cusp of a new era as we explore the complex but promising task of integrating it with artificial intelligence (AI). This fusion has the potential to revolutionize experiment design through automation and streamlined efficiency, thereby reducing manual labor and cognitive load. Our study demonstrates that GPT-4 exhibits impressive proficiency in working with CFs. However, the road to effective integration is filled with significant challenges.

The development of reliable AI interfaces stands as the first hurdle. These interfaces must understand CFs in their entirety and demonstrate proficiency in various tasks such as designing CFs, compiling them into robot commands or human-readable instructions, and even locating CFs using intricate queries. The interfaces should also be capable of querying the sequences tied to the experiments and maintain an inventory awareness. They need to understand and invoke a myriad of API functions related to CFs. This requires an in-depth understanding of the CF specification and the biological processes it encapsulates, coupled with the capacity to handle complex data structures and large sequence files, which constitute substantial computational challenges.

To address the issue of token limits, we propose a decoupling strategy. By assigning unique names to well-defined objects, we can create loosely-coupled references, significantly reducing token usage. This not only simplifies data manipulation across various levels of abstraction but also allows the AI to focus on task-specific requirements within the scope of relevant information.

In light of synthetic biology’s extensive functional scope, we recommend adopting a dynamic plugin system. This would enable the AI to access a wide-ranging function library dynamically, choosing the right function along with its API information for precise execution. This strategy circumvents the need for an AI to be pre-trained on extensive API data and allows for the addition of further functions without necessitating comprehensive AI rewrites.

While it’s crucial to ensure that CFs, the associated sequences, and compiled instructions are stored persistently and readily accessible, the inherently error-prone and fleeting nature of AI memory requires this storage to take place on the plugin side of the interface. The AI should reference these stored objects by their unique names. However, this approach does present challenges, including ensuring that the AI is aware of the objects stored within the plugin and defining how new objects are added and persisted.

In the integration of AI with CFs, biosecurity remains a paramount concern. It is critical to have human oversight to prevent any direct execution of code, particularly when it involves robotic genetic engineering processes. The AI needs to be semantically aware of its tasks and carry out continuous checks against known biohazards such as toxins, virulence factors, and gene drives. As we strive to overcome computational and biosecurity challenges in the integration of AI with CFs, we recognize that the interplay of AI capabilities and synthetic biology, despite its hurdles, holds the key to a future where efficiency, precision, and safety transform the landscape of biological experimentation and discovery.

## Supporting information

All supplemental files

## Supporting Information

Files provided in CF_Supplement.zip:

cf_JSDoc_specification.md: Specification document for JSON representation of CF. cf_shorthand_specification.md: Specification document for CF Shorthand

### Examples Folder

Construction_pSB1A2-Bca9128.txt. CF Shorthand. Example from figure 1 and from OpenWetWare tutorial. Uses PCR, cut and ligate to make a BioBrick. Runs with both PyDNA_CF_Simulator and ConstructionFileSimulator. Will not run with C6-Tools.

Construction_pSB1A2-Bca9128.json. CF JSON. Same experiment as above.

Construction_pSB1A2-Bca9128.xml. CF XML. Same experiment as above.

Construction_pTarg1.txt. CF Shorthand. Replacing the antibiotic marker in pTargetF with ampicillin resistance using two-part PCR-based Golden Gate assembly. Will not work on PyDNA_CF_Simulator, but works with ConstructionFileSimulator.

Construction_TPjoin.txt: CF Shorthand. Plasmid-based Golden Gate of two plasmids with BseRI. Will not work on PyDNA_CF_Simulator, but works with ConstructionFileSimulator.

Construction_pTarg2.txt: CF Shorthand. Replacing the antibiotic marker in pTargetF with a two-part PCR-based Gibson assembly. This also demonstrates usage of optional PCR product size parameters as well as the usage of ‘oligo’ and ‘plasmid’ keywords for sequences. Runs with both PyDNA_CF_Simulator and ConstructionFileSimulator.

Construction_pTarget-tyrB1.txt: CF Shorthand. Replacing the protospacer in pTargetF with a new sequence using SpeI-based EIPCR (cut and ligation of a single PCR product). Runs with both PyDNA_CF_Simulator and ConstructionFileSimulator.

Construction_pBca1100-Bca1111.txt: CF Shorthand. Derived from OpenWetWare tutorial. This demonstrates two capabilities of ConstructionFileSimulator but not implemented in PyDNA_CF_Simulator: SOEing as well as PCR with oligos containing a mismatch in the annealing region.

Construction_pAC-tRNA_N8_Library.txt: CF Shorthand. Annealing and extension (Klenow Extension) of degenerate oligonucleotides, followed by cut and paste. Includes a 3’ enzyme, PstI. Will not work on PyDNA_CF_Simulator, but works with ConstructionFileSimulator.

Construction_PCA.txt: CF Shorthand. A synthetic example of Polymerase Chain Assembly (not tested in the wetlab). Will not work on PyDNA_CF_Simulator, but works with ConstructionFileSimulator.

Lycopene2: Experiment Folder. Insertion of the AtIPI gene into a lycopene production plasmid with 6 different truncations of adjacent sequences. Runs with ConstructionFileSimulator from the command line.

pTP2_reporter: Experiment Folder. A chain of three sequentially executed Construction Files. Runs with ConstructionFileSimulator from the command line.

### Movies Folder

pydna_plugin_test.mov: Demonstration of simulating a PCR reaction via the PyDNA_CF_Simulator ChatGPT plugin.

cfs_experiment.mov: Demonstration of using ConstructionFileSimulator to simulate the pTP2_reporter example.

### Chats Folder

syntax_conversions.html: ChatGPT conversion of CF shorthand to JSON and XML formats. Shared chat: https://chat.openai.com/share/2df31c09-5d2a-4f2b-a264-2b268ce951f6.

cf_to_pydna.html: ChatGPT attempts to convert a CF to a pydna script. Shared chat: https://chat.openai.com/share/cce4f72e-518d-4485-879a-ff98a9ec16b4. Testing of the resulting script: https://colab.research.google.com/drive/1RLDmqEBVl0SNf4lDuGh9x3c22E04V4r2.

design_biobrick.html: ChatGPT attempts to design a BioBrick prefix insertion CF. Shared chat: https://chat.openai.com/share/b618df5f-c319-4a0f-80be-96485a2f33c4.

technician_instructions.html: ChatGPT generates technician protocols from a CF. Shared chat: https://chat.openai.com/share/694a76be-c604-46fc-8872-2c40125d2e00.

autoprotocol_instructions.html: ChatGPT attempts to write an Autoprotocol from a CF. Shared chat: https://chat.openai.com/share/7379b4a3-6c01-489b-a67b-5ae01028f58e.

invasin_native_text.txt: Text from ref. 14 used as a ChatGPT parsing test.

invasin_parse_test.html: ChatGPT parses a CF from scientific journal text. Shared chat: https://chat.openai.com/share/dd951f5b-00e9-42d8-85d2-8fd811f1ab1b.

invasin_cf.txt: CF Shorthand. CF parsed by ChatGPT with sequences added.

invasin_simulation.html: ChatGPT attempts to simulate the invasin CF with no plugin. Shared chat: https://chat.openai.com/share/f1289f66-d4c5-483b-b07c-26071af36a4a.

pydna_plugin_test.html: ChatGPT simulates a PCR reaction using the PyDNA_CF_Simulator plugin. Same as movie. Shared chat: https://chat.openai.com/share/2af29a1f-590c-484f-b48d-7d5ac57fd786.

## Acknowledgements

Evan Cory, Kristen Delgado, Cole Ingamells, Madhumita Kannan, Lauren Mathis, Sisi Morris-Gavrieli, Mona Zheng, students of iGEM at Berkeley and BioE 140L contributed to testing ConstructionFileSimulator and C6-Tools. Though ConstructionFileSimulator predates ChatGPT, all other aspects of this study were done with aid from the AI.

