## Supplementary material for "Enabling AI in Synthetic Biology through Construction File Specification": All supplemental files: autoprotocol_instructions.html

CF to Autoprotocol - A ShareGPT conversation 

Model: GPT-4

### CF Shorthand Specification
#### Layer 0: Basic Concept
A Construction File (CF) is a list of Steps, with each Step representing a specific operation in a molecular biology experiment. Each Step is written on a new line. Parameters are separate by whitespace, preferably TSV. The Step includes the names of input DNA sequence(s), non-sequence parameters, and ends with the name of the product DNA sequence. The input sequences can refer to products of previous steps. The product is the output of the operation.
In addition, a CF can include sequences in the form of 'name sequence' lines.
Comment lines follow '#', '//', or '/\*comment\*/' syntax.
#### Layer 1: Core Operations
This layer defines a core set of operations: PCR, Digest, Ligate, GoldenGate, Gibson, and Transform. Each operation has specific parameters and (optionally) a product name.
#### Examples
- `PCR ForwardPrimer ReversePrimer Template ProductName`
- ForwardPrimer, ReversePrimer, Template: Names of DNA sequences
- `Digest DNA Enzymes FragmentSelection ProductName`
- DNA: Name of the DNA sequence to be digested
- Enzymes: List of enzymes to be used, e.g. EcoRI,BamHI
- FragmentSelection: Index indicating the chosen fragment post-digestion, according to orientation and origin given in the input DNA.
- `Ligate Fragment1 Fragment2 ProductName`
- Fragment1, Fragment2: Names of DNA sequences
- `GoldenGate Fragment1 Fragment2 Enzyme ProductName`
- Fragment1, Fragment2: Names of DNA sequences
- Enzyme: Type IIS-like enzyme to be used along with ligase. e.g. BsaI
- `Gibson Fragment1 Fragment2 ProductName`
- Fragment1, Fragment2: Names of DNA sequences
- `Transform Plasmid Host Antibiotic Temperature ProductName`
- Plasmid: Name of the DNA sequence
- Host: Bacterial strain
- Antibiotics: Antibiotics used, e.g. Amp,Kan
- Temperature: Incubation temperature in Celsius, e.g. 37
Here is an example I wish to discuss:
//Figure 1 Example: PCR, cut, and ligate with palindromic type II six-cutters of two fragments
//Derived from https://openwetware.org/wiki/Arking:JCAOligoTutorial1
### Cloning Steps
PCR ca1067F ca1067R pSB1AK3-b0015 pcrpdt
Digest pcrpdt EcoRI,SpeI 1 pcrdig
Digest pSB1A2-I13521 EcoRI,SpeI 1 vectdig
Ligate pcrdig vectdig lig
Transform lig DH10B Amp 37 pSB1A2-Bca9128

This is a nice shorthand notation to codify molecular biology protocols in an easy-to-read format. Here's what I understand from your Construction File (CF):

The Construction File begins with comments explaining the experiment. The `//` syntax denotes these are comment lines, so they don't perform any actual operations but serve to clarify the protocol or add any necessary information.

This file contains five main steps.

1. `PCR ca1067F ca1067R pSB1AK3-b0015 pcrpdt`

   - This step is a PCR (Polymerase Chain Reaction) amplification using primers `ca1067F` and `ca1067R` on the template DNA sequence `pSB1AK3-b0015`. The PCR product is named `pcrpdt`.
2. `Digest pcrpdt EcoRI,SpeI 1 pcrdig`

   - The PCR product `pcrpdt` is then digested with the enzymes `EcoRI` and `SpeI`. The first fragment resulting from this digestion is selected, as indicated by the '1'. The digested product is named `pcrdig`.
3. `Digest pSB1A2-I13521 EcoRI,SpeI 1 vectdig`

   - In a parallel step, another DNA sequence `pSB1A2-I13521` is also digested with `EcoRI` and `SpeI`. Again, the first fragment is selected and named `vectdig`.
4. `Ligate pcrdig vectdig lig`

   - The digested PCR product (`pcrdig`) and the digested vector (`vectdig`) are ligated together to form a new DNA sequence named `lig`.
5. `Transform lig DH10B Amp 37 pSB1A2-Bca9128`

   - The ligation product `lig` is transformed into the DH10B strain of bacteria. The transformation process uses Ampicillin (Amp) as the antibiotic and the bacteria are incubated at a temperature of 37°C. The resultant transformed bacteria with the new plasmid are named `pSB1A2-Bca9128`.

This way, the process of creating a new DNA sequence via PCR, digestion, ligation, and transformation is neatly summarized in five lines. Each step also generates intermediate products that can be referenced in subsequent steps.

This is from https://autoprotocol.org/specification/, I'll give it to you in installments:
Background
Motivation
There is a crisis of productivity and reproducibility in the life sciences today. Projects that should take weeks end up taking months and the vast majority of published literature struggles to be replicated by independent labs later.
Experimental protocols written in natural language are often ambiguous. For example, the phrases "spin down briefly" and "mix gently" are frequently used in many common protocols and convey much less information than is necessary for operators to reproduce each others' work.
Design Goals
Flexible
Autoprotocol allows for a plethora of possible protocols built from a small set of instructions. No biological knowledge is included in the specification. Adding new instructions is straightforward.
Composable
High levels of complexity are enabled by building up from smaller pieces. It should be possible to start from simple, rock-solid modules and compose them into cutting edge science.
Synthesizable
Autoprotocol is mappable directly to hardware commands for robotic automation. Human interpretation must not be necessary.
Platform Independent
Autoprotocol should be able to be generated and consumed by software written in any language on any platform.
Just Data
Encoded protocols are a linear series of instructions to execute and contain no branching logic or looping constructs evaluatable at runtime.
Learnable
A central design goal of Autoprotocol is the ability for users to extrapolate about how functionality they haven't yet used might work based on the parts they already know and frequently guess correctly.
Conventions In This Document
The key words "MUST", "MUST NOT", "REQUIRED", "SHALL", "SHALL NOT", "SHOULD", "SHOULD NOT", "RECOMMENDED", "MAY", and "OPTIONAL" in this document are to be interpreted as described in IETF RFC 2119.
Syntax
In the code excerpts and examples here, there are a few conventions to know. An unquoted string as a value is a type (for example, the Volume and Time type designations in the "volume" and "duration" fields at right). A quoted string is a literal, as in the "op" value at right. Square brackets denote an array, as in the array of objects.
{
"op": "my\_instr",
"objects": [Container],
"volume": Volume,
"duration": Time,
"count": Int
}
Dimensioned values ("quantities")
Both the volume and duration are quantities, which are strings of the format "magnitude:unit". Duration strings might be 50:second, 12:minute, 50:millisecond, and so on. Similarly, volume strings might be 25:microliter or 5:milliliter. Measures may contain decimals, as in 25.2:microliter. Dimensions are always written singular.

This excerpt is the initial introduction to the AutoProtocol standard, which is a way of formally specifying biological experiments to improve reproducibility and efficiency.

The text covers the motivation behind AutoProtocol, its design goals, the syntax, and the specific conventions followed in the document. The motivation comes from a need for more reproducibility and speed in life science research, where natural language protocols often lead to ambiguity and confusion.

The design goals of AutoProtocol are:

1. **Flexible**: The specification is designed to allow a wide variety of protocols built from a basic set of instructions. It is also easy to add new instructions.
2. **Composable**: The protocol allows complex operations to be built up from smaller modules, allowing the design of cutting-edge science from simple foundations.
3. **Synthesizable**: The AutoProtocol can be directly mapped to hardware commands for robotic automation, eliminating the need for human interpretation.
4. **Platform Independent**: The protocol can be generated and consumed by software written in any language on any platform.
5. **Just Data**: The encoded protocols are a linear series of instructions and do not contain any branching or looping constructs that require runtime evaluation.
6. **Learnable**: AutoProtocol is designed such that users can easily understand its functionality and extrapolate how new, unknown functionality might work based on their knowledge of the existing system.

The document also outlines conventions for language used within it, following the IETF RFC 2119 which defines the interpretation of key words like "MUST", "SHOULD", "RECOMMENDED", etc.

In the given syntax, several conventions are followed. For instance, unquoted strings are types, and quoted strings are literals. Square brackets denote an array. Some examples of syntax are provided. Quantities are defined as strings of "magnitude:unit", where magnitude can be a decimal number and unit is always singular. Examples include "50:second", "12:minute", "25:microliter", etc.

Refs and Datarefs
A ref is an alphanumeric string; a string that contains only letters and numbers and no special characters. Refs are simply easy identifier strings to use to refer to a container defined in an access instruction. Similarly, datarefs are alphanumeric strings used to later identify any data generated by the given instruction. Refs and datarefs must be unique within each protocol.
Containers and Wells
Per the Protocol section, containers are referenced using their ref string. Wells are referenced using a slash syntax :ref/:index, like my\_plate/A1.
Serialization
Autoprotocol protocols are serialized using Javascript Object Notation (JSON). This choice is not intrinsically semantic, but it is mandatory for consistency and compatibility. Alternative serializations such as XML, Protocol Buffers or custom formats shall not be used.
Protocols
Structure
A protocol is defined by three segments:
refs
the set of containers that will be used in the protocol
instructions
the list of instructions to be performed
constraints
constraints on how the instructions should be performed
A ref is a short alphanumeric name given to a container to identify it in later instructions. Every container referenced in a protocol must also be given a destiny: either discarded at the end of the protocol, or stored.
A protocol shall not contain any segments not defined here as mandatory.
{
"refs": {
"dye": {
"id": "ct13zjq79whe",
"store": { "where": "ambient" }
},
"water": {
"id": "ct149x8mea3j",
"store": { "where": "ambient" }
},
"samples": {
"id": "ct3b245kx34l",
"discard": true
},
...
}
}
Once you have references to all the objects you want to work with, you can use them in other instructions by referring to the container itself by its ref or to aliquots within the container with the syntax :ref/:index.
In the protocol snippet at right there are three instructions performing the operations:
Distribute 40 μl from well water/0 into each of test/A1, test/A2, and test/A3.
Distribute 5 μl from well dye/0 into each of test/A1, test/A2, and test/A3.
Centrifuge the plate test for 30 seconds at 2000 g.
Take a 600 nm absorbance reading through wells test/A1, test/A2, and test/A3.
{
"refs": { ... },
"instructions": [
{ "op": "pipette",
"groups": [
{ "distribute": {
"from": "water/0",
"to": [
{ "well": "test/A1",
"volume": "40:microliter" },
{ "well": "test/A2",
"volume": "40:microliter" },
{ "well": "test/A3",
"volume": "40:microliter" }
]
} }, { "distribute": {
"from": "dye/0",
"to": [
{ "well": "test/A1",
"volume": "5:microliter" },
{ "well": "test/A2",
"volume": "5:microliter" },
{ "well": "test/A3",
"volume": "5:microliter" },
]
} }
]
}, {
"op": "spin",
"object": "test",
"duration": "30:second",
"acceleration": "2000:g"
}, {
"op": "absorbance",
"object": "test",
"wells": ["A1", "A2", "A3"],
"wavelength": "600:nanometer"
}]
]
}
Aliquot Paths
While a protocol is just data and does not contain logic (e.g., if/then statements), it is common to use a program that does contain logic to dynamically generate a protocol. For example, the layout of wells on a variable number of plates may change depending on the number of samples being operated on, though the series of operations for each sample is the same (it is "scale invariant"). On the surface, this can make it appear complex to compare protocols over time or across different conditions.
The concept of aliquot paths captures the common scientifically-relevant structure across generated protocols that differ in their overall content due to scale. Two protocols are homomorphic if for every ref in one protocol there is one or more similar ref(s) in the second protocol with the same path. Protocol homomorphism is directional: if one protocol contains additional refs not seen in the other whose paths are independent from the paths of the isomorphic refs (the refs do not interact and constitute completely separate "subroutines" within the protocol), the protocols may still be said to be homomorphic in the context of the refs with common paths.
Put more simply, if there are two protocols that perform the same set of conceptual operations on a different number of samples, adding additional operations and samples that have nothing to do with the existing samples doesn't break the idea that the protocols are "similar, just scaled" for the original samples.
Aliquot paths are important because they allow us to compare logical blocks of operations irrespective of how they're physically configured.

This segment of the AutoProtocol specification provides more detailed instructions on various aspects of the protocol:

1. **Refs and Datarefs**: These are alphanumeric strings that serve as identifiers for containers and data respectively within a protocol. Each ref and dataref must be unique within the protocol.
2. **Containers and Wells**: A container is referenced using its ref string, while a well within a container is referenced using the `:ref/:index` format.
3. **Serialization**: Protocols are serialized using JavaScript Object Notation (JSON). This choice is not related to the semantic meaning of the protocol but for consistency and compatibility across platforms.
4. **Protocols**: A protocol consists of three sections: refs, instructions, and constraints. Refs define the containers used in the protocol, instructions are the actions to be performed, and constraints provide specific limitations on how instructions should be executed. Each container is given a destiny, meaning it's either discarded or stored after the protocol's execution.
5. **Aliquot Paths**: This concept allows the comparison of logical blocks of operations irrespective of their physical configuration. Essentially, it allows one to view two protocols as similar if they perform the same set of conceptual operations on a different number of samples. Even if additional operations and samples are added, the protocols may still be considered homomorphic (similar) in the context of the original samples.

The given JSON snippets show examples of how to structure a protocol, how to reference containers and wells, and how to distribute liquids among wells. An example operation of distributing liquid, centrifuging a plate, and taking an absorbance reading is provided, showing how these operations are encoded in JSON following the AutoProtocol specification.

Definitions
Types
Instruction and ref specification use the following common types.
Primitive Types
Type Definition
Boolean true or false
Float a floating point numeric value
Int an integer numeric value
String any sequence of utf-encoded characters bounded with "
Derived Types
Type Example Value Definition
Aliquot "growth\_plate/A1" an Autoprotocol container and a well index delimited with a / represented as a String
Container "growth\_plate" an Autoprotocol container referenced in the refs section of the protocol represented as a String
Quantity e.g. Volume "5:microliters" a magnitude and a unit delimited with a : represented as a String
Compound {"format": "InChI", "value": "InChI=1S/CH4/h1H4"} a chemical compound defined by the String ‘value’ written following the pattern described by the ‘format’
Type Wrappers
Syntax Example Specification Definition
Enum(..) Enum("one, "two") any one of the enclosed values
Option Option either be the enclosed Type or null
Units
Instruction and ref specification use the following units to represent quantities.
Unit Examples
Acceleration meter/second^2, millimeter/second^2
Amount mole, millimole, micromole, nanomole
AmountConcentration mole/liter, millimole/liter, molar, millimolar
Area meter^2
Capacitance farad, picofarad
ElectricPotential volt, millivolt, microvolt, nanovolt
Frequency hertz, kilohertz, rpm
Length meter, millimeter, micrometer, nanometer
Mass gram, milligram, microgram, nanogram
MassConcentration milligram/milliliter, nanogram/microliter
Power watt, milliwatt, microwatt
Pressure pascal, bar, torr
Temperature celsius, kelvin
Time day, hour, minute, second, millisecond
Velocity meter/second, millimeter/second
Volume liter, milliliter, microliter, nanoliter
VolumeAcceleration milliliter/second^2, microliter/second^2
VolumeConcentration milliliter/milliliter, microliter/microliter
VolumeFlow milliliter/second, microliter/second
Fields
Some common fields that are shared across instructions are defined below.
{
"shake\_path": Enum(
"cw\_orbital",
"ccw\_orbital",
"portrait\_linear",
"landscape\_linear",
"cw\_diamond",
"ccw\_diamond",
"portrait\_down\_double\_orbital",
"landscape\_down\_double\_orbital",
"portrait\_up\_double\_orbital",
"landscape\_up\_double\_orbital"
)
}
Refs
container\_refs
Names: The refs field aliases Containers to descriptive Strings called refs.
Origins: The id field is used to specify the unique identifier of an existing Container. The new field is used to specify that this ref does not yet exist and what type of Container it should be. These two fields are mutually exclusive.
Destinies: The discard field indicates whether the ref should be discarded or not. The store field indicates how a ref should be stored. These two fields are mutually exclusive.
Covers The cover field indicates the type of cover a container is initially covered with. If no cover is specified the container is assumed to be uncovered.
{
"refs": {
String: {
"id": Option,
"new": Option,
"store": Option&lt;{
"where": Option
}&gt;,
"discard": Option,
"cover": Option
}
}
}
Instructions
The instructions field of a Protocol is a made up of a list of Instructions. Instructions are encoded as an op which is the instruction name and optionally a series of additional top-level fields to encode how it should be executed.
Following is the set of instructions currently in the Autoprotocol standard.
acoustic\_transfer
Acoustic liquid handling uses acoustics to fly individual droplets from a source container to a destination one. Most acoustic liquid handlers only support a discrete set of droplet\_size and the volume field of each transfer must be a multiple of it. prevalidate\_sources is used to ensure that the source wells contain enough volume to successfully complete the transfer. source\_volume\_limits are used to overwrite vendor-specified defaults for what volumes should pass prevalidation.
{
"op": "acoustic\_transfer",
"droplet\_size": Option,
"prevalidate\_sources": Option,
"groups": [
{
"transfer": [
{
"from": Aliquot,
"to": Aliquot,
"volume": Volume
}
]
}
],
"source\_volume\_limits": Option&lt;{
"min": Option,
"max": Option
}&gt;
}
cover
Containers must be covered or sealed for storage, incubation, and centrifugation operations (among others). Many instructions including liquid handling operations require that a container be uncovered before use. retrieve\_lid indicates that a lid previously saved by a uncover operation with store\_lid should be used.
{
"op": "cover",
"object": Container,
"lid": String,
"retrieve\_lid": Option
}
flow\_cytometry
Flow cytometry optically detects and characterizes particles suspended in a fluid. For each flow\_cytometry instruction, the channel information will be collected for each sample given the collection conditions specified. stop\_criteria are combined based on the condition specified in trigger\_logic; if left unset, the aquisition\_volume will be used.
{
"op": "flow\_cytometry",
"dataref": String,
"samples": [Aliquot],
"lasers": [
{
"excitation": Option,
"power": Option,
"area\_scaling\_factor": Option,
"channels": [
{
"emission\_filter": {
"shortpass": Option,
"longpass": Option,
"channel\_name": String,
},
"detector\_gain": ElectricPotential,
"measurements": Option&lt;{
"area": Option,
"height": Option,
"width": Option
}&gt;,
"trigger\_threshold": Option,
"trigger\_logic": Option
}
]
}
],
"collection\_conditions": {
"acquisition\_volume": Volume,
"flowrate": VolumeFlow,
"stop\_criteria": Option&lt;{
"volume": Option,
"events": Option,
"time": Option<time>
}&gt;,
"wait\_time": Time,
"mix\_cycles": Int,
"mix\_volume": Volume,
"rinse\_cycles": Int
},
"width\_threshold": Option,
"window\_extension": Option,
"remove\_coincident\_events": Option
}
incubate
The incubate instruction stores a sample in an incubator with the appropriate settings for a given duration.
{
"op": "incubate",
"object": Container,
"where": Enum(
"cold\_20",
"cold\_4",
"ambient",
"warm\_37"
),
"duration": Time,
"shaking": Boolean,
"co2\_percent": Option,
"target\_temperature": Option,
"shaking\_params": Option&lt;{
"frequency": Frequency,
"path": Option,
"amplitude": Option
}&gt;
}
</time>

This part of the AutoProtocol specification includes definitions of types, units, fields, refs, and instructions, as well as their usage in the specification.

1. **Types**: There are both primitive and derived types in AutoProtocol. Primitive types include Boolean, Float, Int, and String. Derived types are Aliquot, Container, Quantity, and Compound, which represent more specific information used in the protocol such as a container with a well index, a container reference, a magnitude with a unit, and a chemical compound, respectively.
2. **Units**: A wide variety of units are used in the protocol to represent quantities. These units include acceleration, amount, concentration, area, capacitance, electric potential, frequency, length, mass, power, pressure, temperature, time, velocity, volume, and their respective sub-types.
3. **Fields**: Fields define specific parameters for the instructions. For example, the "shake\_path" field defines how shaking should occur during an incubate instruction.
4. **Refs (Container Refs)**: Refs are used to give aliases to containers. They can denote either an existing container (using an "id") or a new one to be created (using "new"). They also define how a container should be treated after a protocol is completed, either discarded or stored.
5. **Instructions**: These are encoded as an 'op', which represents the instruction name, and additional fields to encode how the operation should be executed. Examples include acoustic\_transfer (for acoustic liquid handling), cover (to cover containers), flow\_cytometry (for optical detection of particles), and incubate (for storing a sample under specific conditions).

Every instruction has its own set of parameters which need to be filled according to the requirements of the instruction and the protocol itself. The parameters can be mandatory or optional, and they specify in detail how the instruction should be performed. For instance, the "incubate" instruction includes parameters like where the incubation should occur, the duration, whether shaking is required, the percentage of CO2, the target temperature, and specific shaking parameters.

liquid\_handle
The liquid\_handle instruction acts as a framework to allow precise control over liquid handling parameters and express a broad range of liquid handling operations.
The liquid\_handle operation is based around transporting volumes of liquid in and out of locations. Each operation is a locations sequence of location with transports sequences specifying the list of volumes. The position of device components may reset between elements of locations. Transports within the same locations use the same consumables (i.e. tips in the case of air\_displacement liquid handling).
{
"op": "liquid\_handle",
"locations": [
{
"location": Option,
"transports": Option&lt;[
{
"volume": Option,
"pump\_override\_volume": Option,
"flowrate": Option&lt;{
"target": Option,
"initial": Option,
"cutoff": Option,
"acceleration": Option&lt;
VolumeAcceleration
&gt;,
"deceleration": Option&lt;
VolumeAcceleration
&gt;
}&gt;,
"delay\_time": Option<time>,
"mode\_params": Option&lt;{
"liquid\_class": Option,
"tip\_position": Option&lt;{
"position\_x": Option&lt;{
"position": Option,
"move\_rate": Option&lt;{
"target": Option,
"acceleration": Option&lt;
Acceleration
&gt;
}&gt;
}&gt;,
"position\_y": Option&lt;{
"position": Option,
"move\_rate": Option&lt;{
"target": Option,
"acceleration": Option&lt;
Acceleration
&gt;
}&gt;
}&gt;,
"position\_z": Option&lt;{
"offset": Option,
"move\_rate": Option&lt;{
"target": Option,
"acceleration": Option&lt;
Acceleration
&gt;
}&gt;,
"reference": Option,
"detection": Option&lt;{
"method": Option,
"threshold": Option,
"duration": Option<time>,
"fallback": Option
}&gt;
}&gt;
}
}&gt;
}
]&gt;,
"temperature": Option,
"density": Option
}
],
"mode": Option
"mode\_params": Option&lt;{
"tip\_type": Option
}&gt;,
"shape": Option&lt;{
"rows": Int,
"columns": Int,
"format": Option
}&gt;
}
measure\_mass
The measure\_mass instruction can be used to determine the mass of a sample (container). The execution is vendor specific and may or may not consume a fraction of the sample. The accuracy of results is vendor specific.
{
"op": "measure\_mass",
"object": Container,
"dataref": String
}
measure\_volume
The measure\_volume instruction can be used to determine the volume of a sample. The execution is vendor specific and may or may not consume a fraction of the sample. The accuracy of results is vendor specific.
{
"op": "measure\_volume",
"object": [Aliquot],
"dataref": String
}
provision
The provision instruction encodes adding some amount of an external resource to an aliquot or series of aliquots.
{
"op": "provision",
"resource\_id": String,
"measurement\_mode": Option
"to": [
Option&lt;{
"well": Aliquot,
"volume": Volume,
"dispense\_velocity": Option,
"mix\_after": Option&lt;{
"volume" Volume,
"repetitions": Int,
"velocity": Option
}&gt;
}&gt;,
Option&lt;{
"well": Aliquot,
"mass": Mass
}&gt;
]
}
seal
Containers must be covered or sealed for storage, incubation, and centrifugation operations (among others). Seal types have useful properties ranging from optical clarity to gas permeability. Seals can be applied by either thermal or adhesive sealers which result in different seal integrity. thermal seals can be applied with a range of temperatures and durations that can be optimized for different plate types. Many instructions including liquid handling operations require that a container be uncovered before use.
{
"op": "seal",
"object": Container,
"type": String,
"mode": Option,
"mode\_params": Option&lt;{
"temperature": Option,
"duration": Option<time>
}&gt;
}
</time></time></time>

Here are more instructions used in the AutoProtocol standard:

1. **liquid\_handle**: The liquid\_handle instruction is used for a wide range of liquid handling operations, allowing precise control over parameters. This instruction involves transporting volumes of liquid in and out of locations, with each operation consisting of a sequence of locations and transport sequences specifying the list of volumes. Each transport can have parameters including volume, flow rate, delay time, and mode parameters.
2. **measure\_mass**: This instruction determines the mass of a sample container. The specifics of the execution and the accuracy of the results depend on the vendor.
3. **measure\_volume**: The measure\_volume instruction is used to determine the volume of a sample. Like measure\_mass, the execution specifics and the result's accuracy are vendor-specific.
4. **provision**: This instruction represents the addition of an external resource to an aliquot or series of aliquots. It can be used to specify the addition of a certain volume or mass of a resource to a specific well.
5. **seal**: The seal instruction is used for covering or sealing containers for storage, incubation, and centrifugation operations among others. Seals can be applied either by thermal or adhesive sealers, which result in different seal integrity.

Each of these instructions has specific parameters that need to be filled according to the requirements of the instruction and the protocol. The parameters can be either mandatory or optional and specify in detail how the instruction should be executed.

spectrophotometry
The spectrophotometry instruction encodes one or a series of plate reading steps executed on a single container with the same device. This could be executed once, or at a defined interval, across some total duration. There are 4 valid modes (absorbance, fluorescence, luminescence, and shake) that each accept a different set of mode\_params
```
{
"op": "spectrophotometry",
"dataref": String,
"object": Container,
"interval": Option<time>,
"num\_intervals": Option,
"temperature": Option,
"shake\_before": Option&lt;{
"duration": Time,
"frequency": Option,
"amplitude": Option,
"path": Option
}&gt;,
"groups": [
Option&lt;{
"mode": "absorbance",
"mode\_params": {
"wells": [Aliquot],
"wavelength": [Length],
"num\_flashes": Option,
"settle\_time": Option<time>,
"read\_position": Option,
"position\_z": Option&lt;{
"manual": Option&lt;{
"displacement": Length,
"reference": Enum(
"plate\_bottom",
"plate\_top",
"well\_bottom",
"well\_top"
)
}&gt;,
"calculated\_from\_wells": Option&lt;{
"wells": [Aliquot],
"heuristic": Enum(
"max\_mean\_read\_without\_saturation",
"closest\_length\_without\_saturation"
)
}&gt;,
}&gt;
}
}&gt;,
Option&lt;{
"mode": "fluorescence",
"mode\_params": {
"excitation": [{
"shortpass": Option,
"longpass": Option,
"ideal": Option
}],
"emission": [{
"shortpass": Option,
"longpass": Option,
"ideal": Option
}],
"num\_flashes": Option,
"settle\_time": Option<time>,
"lag\_time": Option<time>,
"integration\_time": Option<time>,
"gain": Option,
"read\_position": Option,
"position\_z": Option&lt;{
"manual": Option&lt;{
"displacement": Length,
"reference": Enum(
"plate\_bottom",
"plate\_top",
"well\_bottom",
"well\_top"
)
}&gt;,
"calculated\_from\_wells": Option&lt;{
"wells": [Aliquot],
"heuristic": Enum(
"max\_mean\_read\_without\_saturation",
"closest\_length\_without\_saturation"
)
}&gt;,
}&gt;
}
}&gt;,
Option&lt;{
"mode": "luminescence",
"mode\_params": {
"wells": [Aliquot],
"num\_flashes": Option,
"settle\_time": Option<time>,
"integration\_time": Option<time>,
"gain": Option,
"read\_position": Option,
"position\_z": Option&lt;{
"manual": Option&lt;{
"displacement": Length,
"reference": Enum(
"plate\_bottom",
"plate\_top",
"well\_bottom",
"well\_top"
)
}&gt;,
"calculated\_from\_wells": Option&lt;{
"wells": [Aliquot],
"heuristic": Enum(
"max\_mean\_read\_without\_saturation",
"closest\_length\_without\_saturation"
)
}&gt;,
}
}&gt;,
Option&lt;{
"mode": "shake",
"mode\_params": {
"duration": Option<time>,
"frequency": Option,
"amplitude": Option,
"path": Option
}
}&gt;
]
}
```
spin
The spin instruction is used to represent a series of centrifugation steps. The inward and outward flow\_direction encodes spinning the contents into or out of of a container respectively. The operation is repeated with the appropriate direction for each element in spin\_directions.
{
"op": "spin",
"object": Container,
"acceleration": Acceleration,
"duration": Time,
"flow\_direction": Option,
"spin\_direction": [
Enum("cw", "ccw")
]
}
uncover
Containers must be covered or sealed for storage, incubation, and centrifugation operations (among others). Many instructions including liquid handling operations require that a container be uncovered before use. store\_lid indicates that the lid should be saved for some subsequent cover instruction with retrieve\_lid.
{
"op": "uncover",
"object": Container,
"store\_lid": Option
}
unseal
Containers must be covered or sealed for storage, incubation, and centrifugation operations (among others). Many instructions including liquid handling operations require that a container be uncovered before use.
{
"op": "unseal",
"object": Container
}
Constraints
time\_constraints
Time constraints encode a temporal relationship between two time points, from and to.
Each of the time points must specify exactly one of their optional fields. ref\_start and ref\_end encode the points at which a Container leaves its origin and enters its destiny respectively. instruction\_start and instruction\_end encode the points at the beginning and end of an instruction’s execution; the instruction is represented by its 0-indexed position within the instructions list.
Each time constraint may include any combination of the less\_than, more\_than, and ideal fields. less\_than and more\_than constraints encode the minimum and maximum amount of time that is allowable between the two time points. ideal constraints encode the intended timing between two time points as well as the optimization\_cost by which these fields should be weighted.
{
"time\_constraints": Option&lt;[
{
"from": {
"ref\_start": Option,
"ref\_end": Option,
"instruction\_start": Option,
"instruction\_end": Option
},
"to": {
"ref\_start": Option,
"ref\_end": Option,
"instruction\_start": Option,
"instruction\_end": Option
},
"less\_than": Option<time>,
"more\_than": Option<time>,
"ideal": Option&lt;{
"value": Time,
"optimization\_cost": Option
}&gt;
}
]&gt;
}</time></time></time></time></time></time></time></time></time></time>

Let's go through these additional Autoprotocol instructions and their purpose:

1. **spectrophotometry**: This instruction includes one or a series of plate reading steps executed on a single container with the same device. It may be executed once, or at a defined interval, across some total duration. The instruction includes different modes like absorbance, fluorescence, luminescence, and shake, each of which accepts a different set of parameters.
2. **spin**: The spin instruction is used to represent a series of centrifugation steps. The flow direction determines if the spin will force the contents in or out of the container. The operation is repeated with the appropriate direction for each element in spin\_directions.
3. **uncover**: The uncover instruction is used to remove the cover from a container. It's generally used in conjunction with other instructions that require the container to be uncovered before use, such as liquid handling operations. The `store_lid` option indicates whether the lid should be saved for a later cover instruction with `retrieve_lid`.
4. **unseal**: Similar to the uncover instruction, unseal is used to remove a seal from a container. This instruction is typically used before executing instructions that require the container to be unsealed, like liquid handling operations.
5. **Constraints / time\_constraints**: These represent temporal relationships between two time points and specify the minimum and maximum time allowed between these points, as well as an ideal timing and how strictly it should be adhered to. These constraints can be applied to the start and end times of a container's journey or the execution of an instruction.

These additional instructions help in enhancing the level of control and precision that can be achieved in an experiment. They offer options to customize the timing, measurement, and preparation of samples in an automated workflow.

Write an Autoprotocol to perform an experiment defined by this CF:
PCR ca1067F ca1067R pSB1AK3-b0015 pcrpdt
Digest pcrpdt EcoRI,SpeI 1 pcrdig
Digest pSB1A2-I13521 EcoRI,SpeI 1 vectdig
Ligate pcrdig vectdig lig
Transform lig DH10B Amp 37 pSB1A2-Bca9128

Sure, I'll write down a basic Autoprotocol that would represent the experiment you've defined. It's worth noting that the protocol will be very generic as it does not include specific parameters such as volumes, durations, etc.

```
jsonCopy code


{
  "instructions": [
    {
      "op": "pcr",
      "dataref": "ca1067F_ca1067R_pSB1AK3_b0015_pcrpdt",
      "reagents": [
        "ca1067F",
        "ca1067R",
        "pSB1AK3-b0015"
      ],
      "object": "pcrpdt"
    },
    {
      "op": "digest",
      "object": "pcrpdt",
      "enzymes": [
        "EcoRI",
        "SpeI"
      ],
      "digests": 1,
      "dataref": "pcrdig"
    },
    {
      "op": "digest",
      "object": "pSB1A2-I13521",
      "enzymes": [
        "EcoRI",
        "SpeI"
      ],
      "digests": 1,
      "dataref": "vectdig"
    },
    {
      "op": "ligate",
      "fragments": [
        "pcrdig",
        "vectdig"
      ],
      "dataref": "lig"
    },
    {
      "op": "transform",
      "object": "lig",
      "strain": "DH10B",
      "antibiotic": "Amp",
      "temperature": 37,
      "output": "pSB1A2-Bca9128"
    }
  ]
}
```

This protocol includes five main steps:

1. **PCR**: Polymerase chain reaction using the ca1067F and ca1067R primers, and the pSB1AK3-b0015 DNA template. The product of the reaction is referred to as "pcrpdt".
2. **Digest**: Digestion of the PCR product "pcrpdt" using the restriction enzymes EcoRI and SpeI. The product is referred to as "pcrdig".
3. **Digest**: Digestion of the plasmid "pSB1A2-I13521" using the restriction enzymes EcoRI and SpeI. The product is referred to as "vectdig".
4. **Ligate**: Ligation of the two digested fragments "pcrdig" and "vectdig". The product is referred to as "lig".
5. **Transform**: Transformation of the ligated product "lig" into the bacterial strain DH10B, followed by selection on ampicillin at 37 degrees Celsius. The final product is referred to as "pSB1A2-Bca9128".

This protocol is a basic representation of the experiment you described and can be adjusted to fit the specific requirements of your experiment.

That doesn't follow the ontology defined by the spec

I apologize for the misunderstanding. Autoprotocol is a JSON-based protocol format that abstracts specific lab operations. However, the provided experimental outline does not offer enough details to craft a fully specified Autoprotocol. For instance, volumes, concentrations, temperatures, durations, devices, etc., aren't provided in the given plan.

Here's an attempt to sketch a generic Autoprotocol, assuming some parameters:

```
jsonCopy code


{
  "refs": {
    "pcr_plate": {
      "new": "96-pcr",
      "store": {
        "where": "ambient"
      }
    },
    "digestion_plate": {
      "new": "96-pcr",
      "store": {
        "where": "ambient"
      }
    },
    "ligation_plate": {
      "new": "96-pcr",
      "store": {
        "where": "ambient"
      }
    }
  },
  "instructions": [
    {
      "op": "thermocycle",
      "object": "pcr_plate",
      "groups": [
        // Specify PCR conditions here.
      ]
    },
    {
      "op": "digest",
      "reagents": [
        // Specify digest reagents here.
      ],
      "object": "digestion_plate",
      "time": "1:00:00"
    },
    {
      "op": "digest",
      "reagents": [
        // Specify digest reagents here.
      ],
      "object": "digestion_plate",
      "time": "1:00:00"
    },
    {
      "op": "ligate",
      "reagents": [
        // Specify ligation reagents here.
      ],
      "object": "ligation_plate",
      "time": "1:00:00"
    },
    {
      "op": "transform",
      "dataref": "transformation_result",
      "object": "ligation_plate",
      "cell_strain": "DH10B",
      "antibiotic": "Amp",
      "temperature": "37.0:celsius"
    }
  ]
}
```

In this protocol, I'm assuming that we are using 96-well plates ("96-pcr") for all the operations. PCR, digestion, and ligation are happening in different plates, and the "object" of each operation is a reference to these plates. For each operation, you would need to specify the exact parameters for your experiment.

Please adjust the above protocol to better fit your needs or provide additional experiment details for a more precise protocol.
