## Supplementary material for "Enabling AI in Synthetic Biology through Construction File Specification": All supplemental files: cf_to_pydna.html

The example you provided demonstrates a complete molecular biology experiment in a Construction File (CF) format. It can be dissected step-by-step:

1. `PCR ca1067F ca1067R pSB1AK3-b0015 pcrpdt`: This line means a PCR reaction is carried out using primers `ca1067F` and `ca1067R` on the template DNA sequence `pSB1AK3-b0015`. The resultant product is named `pcrpdt`.
2. `Digest pcrpdt EcoRI,SpeI 1 pcrdig`: This command indicates a DNA digestion step where the DNA product from the previous PCR step (`pcrpdt`) is digested by the enzymes `EcoRI` and `SpeI`. The first fragment (as specified by `1`) resulting from this digestion is selected and named `pcrdig`.
3. `Digest pSB1A2-I13521 EcoRI,SpeI 1 vectdig`: This step is similar to the previous one but performed on a different DNA sequence named `pSB1A2-I13521`. The selected fragment is named `vectdig`.
4. `Ligate pcrdig vectdig lig`: This command represents a ligation process where the two fragments `pcrdig` and `vectdig` from the previous digestion steps are ligated together. The product from this ligation step is named `lig`.
5. `Transform lig DH10B Amp 37 pSB1A2-Bca9128`: The last command is the transformation of the plasmid `lig` into `DH10B` bacterial strain. The transformation culture is then subjected to antibiotic selection (`Amp`) at `37` degrees Celsius. The resulting transformed strain is named `pSB1A2-Bca9128`.

This CF format serves as a precise and succinct way to outline molecular biology experiments, allowing for clear communication and interpretation of experimental design and steps.

Write a pydna script I can run in Google Colab to simulate this CF

Sure, here's a Python script using the `pydna` library that you can run in Google Colab to simulate the Construction File (CF) you've provided.

Note that the actual sequences for primers, templates, and plasmids aren't provided, so I've added placeholders. In your actual script, you would replace these with actual DNA sequences.

```
pythonCopy code


# Import necessary libraries
from pydna.dseq import Dseq
from pydna.amplify import pcr
from pydna.assembly import Assembly
from pydna.readers import read
from pydna.genbank import Genbank

# Install Biopython if not done already
!pip install biopython pydna

# Connect to Genbank
gb = Genbank("")  # Replace with your email

# Define sequences (replace "ATG..." with actual sequences)
ca1067F = Dseq("ATG...")
ca1067R = Dseq("ATG...")
pSB1AK3_b0015 = gb.nucleotide("Part:BBa_B0015")  # Fetch sequence from Genbank

# PCR Step
pcrpdt = pcr(ca1067F, ca1067R, pSB1AK3_b0015)

# Define enzymes
from Bio.Restriction import EcoRI, SpeI
enzymes = [EcoRI, SpeI]

# Digest Steps
pcrdig = pcrpdt.cut(enzymes)[0]  # Assuming fragment of interest is the first one

pSB1A2_I13521 = gb.nucleotide("Part:BBa_I13521")  # Fetch sequence from Genbank
vectdig = pSB1A2_I13521.cut(enzymes)[0]  # Assuming fragment of interest is the first one

# Ligation Step
assembly = Assembly([pcrdig, vectdig])
lig = assembly.circular_products()[0]

# Write output sequences to files
with open("pcrpdt.gb", "w+") as file:
    file.write(pcrpdt.format("genbank"))

with open("lig.gb", "w+") as file:
    file.write(lig.format("genbank"))
```

This script provides a simulation for the steps in the CF but doesn't fully cover the `Transform` operation as the biological transformation process is not straightforward to simulate in a script (i.e., it involves biological processes like transformation efficiency, antibiotic selection, etc., that cannot be accurately modeled). However, the output files `pcrpdt.gb` and `lig.gb` are the desired end products of your genetic manipulation, representing the result of transformation.
