## Supplementary material for "Enabling AI in Synthetic Biology through Construction File Specification": All supplemental files: design_biobrick.html

Ok, I understand the specifications for the Construction File (CF) shorthand notation you provided. It seems to be a structured method for specifying steps and parameters in molecular biology experiments, using different operations such as PCR, Digest, Ligate, GoldenGate, Gibson, and Transform.

pLefty is a BioBrick plasmid encoding the Ptet promoter part flanked by the standard RFC10 polylinker. pRighty is similarly a BioBrick plasmid with an rbs.GFP.terminator cassette. Write a construction file for making a rbs.GFP.terminator part-containing plasmid using a prefix insertion reaction.

Based on your description, here's a basic Construction File (CF) for the operation. I'm assuming you would digest both pLefty and pRighty with EcoRI and PstI, which are commonly used in BioBrick construction (RFC10). The product of this operation will be a new plasmid that contains the rbs.GFP.terminator part.

```
graphqlCopy code


# This is the CF for making a rbs.GFP.terminator part-containing plasmid

# Step 1: Digest pLefty with EcoRI and PstI to create Fragment1
Digest pLefty EcoRI,PstI 1 Fragment1

# Step 2: Digest pRighty with EcoRI and PstI to create Fragment2
Digest pRighty EcoRI,PstI 1 Fragment2

# Step 3: Ligate Fragment1 and Fragment2 to create the new plasmid
Ligate Fragment1 Fragment2 NewPlasmid
```

Please make sure to adjust the `FragmentSelection` parameter if the fragments after digestion are not the first one.

Not quite right. Here is some documentation:
Loading menubar.....
Assembly:Standard assembly
Help:Standard Assembly (zoom)
BioBricks have been designed to be assembled using normal cloning techniques. Two BioBrick parts, for example, one blue and one green, can be assembled into a blue-green system by a process called BioBrick Standard Assembly, illustrated at the left.
The BioBrick is flanked by restriction sites, comprised of the BioBrick Prefix and Suffix, between the EcoRI and XbaI cutting sites on the left and SpeI and PstI on the right. For the assembly shown, the blue part is cut out of its plasmid with the enzymes EcoRI and SpeI. The resulting fragment is called the insert because it will be inserted into the plasmid containing the other part.
In a separate reaction, a gap is cut in the plasmid containing the green part using EcoRI and XbaI. Using gel electrophoresis, the insert for the blue part and the cut plasmid containing the green part are purified and the unwanted fragments discarded. The purified insert and cut plasmid are mixed under the right conditions to allow the E sticky ends to come together and the S sticky ends to come together with the X sticky ends. Once this happens, the DNA backbone is re-ligated and the resulting blue-green part can be transformed into E.coli cells. These cells may be grown to produce as much of blue.green part as desired.
Each cycle of BioBrick assembly joins two parts together and in theory should only take a few hours to perform. However, less experienced assemblers may find the procedure takes a couple of days to a week. Sometimes, the assembly process fails to produce the right assembly or even to produce anything. Such results may come about because the product is not stable in E.coli and gets rejected or mutated by the cells.
An alternative to BioBrick Standard Assembly is Three-Antibiotic (3A) Assembly.

I apologize for the misunderstanding in the previous response. I appreciate you providing the additional information about the standard BioBrick assembly process.

From your description, it appears you're using the EcoRI and SpeI enzymes to cut out the insert from pLefty, while using EcoRI and XbaI enzymes to create a gap in the pRighty plasmid. Subsequently, these two are ligated together and transformed into E.coli cells.

Here's the corrected Construction File:

```
plaintextCopy code


# This is the CF for making a rbs.GFP.terminator part-containing plasmid

# Step 1: Digest pLefty with EcoRI and SpeI to create Fragment1
Digest pLefty EcoRI,SpeI 1 Fragment1

# Step 2: Digest pRighty with EcoRI and XbaI to create the plasmid backbone
Digest pRighty EcoRI,XbaI 1 PlasmidBackbone

# Step 3: Ligate Fragment1 and PlasmidBackbone to create the new plasmid
Ligate Fragment1 PlasmidBackbone NewPlasmid

# Step 4: Transform the NewPlasmid into E.coli
Transform NewPlasmid E.coli Amp 37 NewPlasmidInEColi
```

As before, please adjust the `FragmentSelection` parameter if the fragment of interest is not the first one in the electrophoresis results. Also, ensure to use the correct antibiotics and incubation temperature according to your laboratory standards in the `Transform` step.

In determining the fragmentSelection index, you can assume the plasmids are oriented as beginning with the 5' sequencing site, then the EcoRI site, then the XbaI site and part, then SpeI, then the rest of the plasmid. Thus, the backbone fragment would be index 0 and the part fragment index 1.

Thank you for that additional information. Taking that into account, here is the corrected Construction File:

```
plaintextCopy code


# This is the CF for making a rbs.GFP.terminator part-containing plasmid

# Step 1: Digest pLefty with EcoRI and SpeI to create Fragment1
Digest pLefty EcoRI,SpeI 1 Fragment1

# Step 2: Digest pRighty with EcoRI and XbaI to create the plasmid backbone
Digest pRighty EcoRI,XbaI 0 PlasmidBackbone

# Step 3: Ligate Fragment1 and PlasmidBackbone to create the new plasmid
Ligate Fragment1 PlasmidBackbone NewPlasmid

# Step 4: Transform the NewPlasmid into E.coli
Transform NewPlasmid E.coli Amp 37 NewPlasmidInEColi
```

Now, the `FragmentSelection` parameter in the Digest step refers to the correct fragments according to the orientation you specified. In the Digest operation for pLefty, the desired fragment is the second one (index 1), which is the insert part. In the Digest operation for pRighty, the desired fragment is the first one (index 0), which is the plasmid backbone.
