## Supplementary material for "Enabling AI in Synthetic Biology through Construction File Specification": All supplemental files: invasin_parse_test.html

CF Shorthand Specification - A ShareGPT conversation 

Model: GPT-4

### CF Shorthand Specification
#### Layer 0: Basic Concept
A Construction File (CF) is a list of Steps, with each Step representing a specific operation in a molecular biology experiment. Each Step is written on a new line. Parameters are separate by whitespace, preferably TSV. The Step includes the names of input DNA sequence(s), non-sequence parameters, and ends with the name of the product DNA sequence. The input sequences can refer to products of previous steps. The product is the output of the operation.
In addition, a CF can include sequences in the form of 'name sequence' lines.
Comment lines follow '#', '//', or '/\*comment\*/' syntax.
#### Layer 1: Core Operations
This layer defines a core set of operations: PCR, Digest, Ligate, GoldenGate, Gibson, and Transform. Each operation has specific parameters and (optionally) a product name.
#### Examples
- `PCR ForwardPrimer ReversePrimer Template ProductName`
- ForwardPrimer, ReversePrimer, Template: Names of DNA sequences
- `Digest DNA Enzymes FragmentSelection ProductName`
- DNA: Name of the DNA sequence to be digested
- Enzymes: List of enzymes to be used, e.g. EcoRI,BamHI
- FragmentSelection: Index indicating the chosen fragment post-digestion, according to orientation and origin given in the input DNA.
- `Ligate Fragment1 Fragment2 ProductName`
- Fragment1, Fragment2: Names of DNA sequences
- `GoldenGate Fragment1 Fragment2 Enzyme ProductName`
- Fragment1, Fragment2: Names of DNA sequences
- Enzyme: Type IIS-like enzyme to be used along with ligase. e.g. BsaI
- `Gibson Fragment1 Fragment2 ProductName`
- Fragment1, Fragment2: Names of DNA sequences
- `Transform Plasmid Host Antibiotic Temperature ProductName`
- Plasmid: Name of the DNA sequence
- Host: Bacterial strain
- Antibiotics: Antibiotics used, e.g. Amp,Kan
- Temperature: Incubation temperature in Celsius, e.g. 37
Confirm you understand with "ok"

Ok, I understand. You're defining a shorthand specification for a Construction File (CF) that outlines the steps in a molecular biology experiment. The CF includes a list of Steps, each representing a specific operation and written on a new line. Each step involves input DNA sequences, certain parameters, and a product name.

The six core operations are PCR, Digest, Ligate, GoldenGate, Gibson, and Transform, each with its own specific parameters and an optional product name.

Comments can be included using '#' or '//', or by enclosing them in '/\* \*/'. Sequences can also be included in the format 'name sequence'.

Interpret the following as CF. Assume all the input materials are oligonucleotides or plasmid DNAs. Include any steps that are implied but unstated, but stay within the scope of the facts provided.
Invasin was PCR-amplified from pAC-TetInv with oligonucleotides ca877F (5'-GAGTTGGATCCNNNGGAGNNNNNNRTGNTGGTTTTCCAGCCAATCAGTG-3') and ca606R (5'-GTCGACGGCGCTATTCAGATCCTC-3'), digested with BamHI and EcoRI, and inserted in similar sites of pBACr-Fdh and pBACr-Ara affording ribosome binding site libraries.

Sure, based on the information provided, here's an interpretation of this process into the Construction File (CF) format. Please note that the exact interpretation might vary depending on additional context and detailed procedures, which are not provided here.

First, I will list the sequences:

```
bashCopy code


#Sequence information
pAC-TetInv pAC-TetInvSequence
ca877F GAGTTGGATCCNNNGGAGNNNNNNRTGNTGGTTTTCCAGCCAATCAGTG
ca606R GTCGACGGCGCTATTCAGATCCTC
pBACr-Fdh pBACr-FdhSequence
pBACr-Ara pBACr-AraSequence
```

Then, the steps of the process:

```
bashCopy code


# Steps
PCR ca877F ca606R pAC-TetInv InvasinPCR
Digest InvasinPCR BamHI,EcoRI 1 InvasinFragment
Digest pBACr-Fdh BamHI,EcoRI 1 pBACr-FdhFragment
Digest pBACr-Ara BamHI,EcoRI 1 pBACr-AraFragment
Ligate InvasinFragment pBACr-FdhFragment pBACr-FdhInvasin
Ligate InvasinFragment pBACr-AraFragment pBACr-AraInvasin
```

In the steps above:

1. PCR is carried out with ca877F and ca606R oligonucleotides on pAC-TetInv to amplify the Invasin gene (named as 'InvasinPCR').
2. The product 'InvasinPCR' is then digested with BamHI and EcoRI enzymes to produce a fragment 'InvasinFragment'.
3. Similarly, 'pBACr-Fdh' and 'pBACr-Ara' plasmids are digested with the same enzymes to produce fragments 'pBACr-FdhFragment' and 'pBACr-AraFragment'.
4. Then, the 'InvasinFragment' is ligated with the 'pBACr-FdhFragment' to form 'pBACr-FdhInvasin'.
5. In a parallel ligation, 'InvasinFragment' is also ligated with the 'pBACr-AraFragment' to form 'pBACr-AraInvasin'.

Please note that the selection of the correct fragment after digestion ('1' in the Digest steps) might need to be adjusted based on the actual digestion result. And the precise sequences of the plasmids ('pAC-TetInvSequence', 'pBACr-FdhSequence', 'pBACr-AraSequence') should be replaced with the actual sequences.
