## Supplementary material for "Enabling AI in Synthetic Biology through Construction File Specification": All supplemental files: invasin_simulation.html

CF Shorthand Specification - A ShareGPT conversation 

Model: GPT-4

### CF Shorthand Specification
#### Layer 0: Basic Concept
A Construction File (CF) is a list of Steps, with each Step representing a specific operation in a molecular biology experiment. Each Step is written on a new line. Parameters are separate by whitespace, preferably TSV. The Step includes the names of input DNA sequence(s), non-sequence parameters, and ends with the name of the product DNA sequence. The input sequences can refer to products of previous steps. The product is the output of the operation.
In addition, a CF can include sequences in the form of 'name sequence' lines.
Comment lines follow '#', '//', or '/\*comment\*/' syntax.
#### Layer 1: Core Operations
This layer defines a core set of operations: PCR, Digest, Ligate, GoldenGate, Gibson, and Transform. Each operation has specific parameters and (optionally) a product name.
#### Examples
- `PCR ForwardPrimer ReversePrimer Template ProductName`
- ForwardPrimer, ReversePrimer, Template: Names of DNA sequences
- `Digest DNA Enzymes FragmentSelection ProductName`
- DNA: Name of the DNA sequence to be digested
- Enzymes: List of enzymes to be used, e.g. EcoRI,BamHI
- FragmentSelection: Index indicating the chosen fragment post-digestion, according to orientation and origin given in the input DNA.
- `Ligate Fragment1 Fragment2 ProductName`
- Fragment1, Fragment2: Names of DNA sequences
- `GoldenGate Fragment1 Fragment2 Enzyme ProductName`
- Fragment1, Fragment2: Names of DNA sequences
- Enzyme: Type IIS-like enzyme to be used along with ligase. e.g. BsaI
- `Gibson Fragment1 Fragment2 ProductName`
- Fragment1, Fragment2: Names of DNA sequences
- `Transform Plasmid Host Antibiotic Temperature ProductName`
- Plasmid: Name of the DNA sequence
- Host: Bacterial strain
- Antibiotics: Antibiotics used, e.g. Amp,Kan
- Temperature: Incubation temperature in Celsius, e.g. 37

### CF Shorthand Specification

#### Layer 0: Basic Concept

A Construction File (CF) is a text document that describes a series of steps in a molecular biology experiment. Each line represents a unique step, and these steps are performed in the order they appear in the file. Each line includes the names of input DNA sequences, any necessary non-sequence parameters, and the name of the resulting product DNA sequence. Input sequences can be the products of previous steps, thus establishing a workflow.

In addition to steps, a CF can include DNA sequences, which are written as 'name sequence' lines.

Comments can be added using '#', '//' or '/\* \*/' syntax, and these will not affect the operations described in the CF.

#### Layer 1: Core Operations

There are six core operations, each with their own set of parameters:

1. PCR: Polymerase Chain Reaction, a method used to create copies of specific DNA segments.

   Syntax: `PCR ForwardPrimer ReversePrimer Template ProductName`
2. Digest: The process of cutting DNA at specific sites using restriction enzymes.

   Syntax: `Digest DNA Enzymes FragmentSelection ProductName`
3. Ligate: The action of joining two DNA molecules together.

   Syntax: `Ligate Fragment1 Fragment2 ProductName`
4. GoldenGate: A type of DNA assembly that uses type IIS restriction enzymes and T4 DNA ligase.

   Syntax: `GoldenGate Fragment1 Fragment2 Enzyme ProductName`
5. Gibson: A method for assembling small DNA fragments, especially useful when making larger constructs like genes, pathways, plasmids, and genomes.

   Syntax: `Gibson Fragment1 Fragment2 ProductName`
6. Transform: The process of introducing foreign DNA into a cell.

   Syntax: `Transform Plasmid Host Antibiotic Temperature ProductName`

#### Examples

1. PCR: To amplify a specific DNA segment, the CF line might look like:

   `PCR p1 p2 DNA1 DNA2`

   where 'p1' and 'p2' are the forward and reverse primers, 'DNA1' is the template DNA sequence, and 'DNA2' is the product DNA sequence.
2. Digest: To digest a DNA sequence, the CF line might look like:

   `Digest DNA1 EcoRI,BamHI 2 DNA2`

   where 'DNA1' is the input DNA sequence, 'EcoRI,BamHI' are the restriction enzymes to be used, '2' is the fragment selection index, and 'DNA2' is the product DNA sequence.
3. Ligate: To ligate two DNA fragments, the CF line might look like:

   `Ligate DNA1 DNA2 DNA3`

   where 'DNA1' and 'DNA2' are the input DNA sequences, and 'DNA3' is the product DNA sequence.
4. GoldenGate: To assemble DNA fragments using the Golden Gate method, the CF line might look like:

   `GoldenGate DNA1 DNA2 BsaI DNA3`

   where 'DNA1' and 'DNA2' are the input DNA sequences, 'BsaI' is the type IIS-like enzyme to be used, and 'DNA3' is the product DNA sequence.
5. Gibson: To assemble DNA fragments using the Gibson method, the CF line might look like:

   `Gibson DNA1 DNA2 DNA3`

   where 'DNA1' and 'DNA2' are the input DNA sequences, and 'DNA3' is the product DNA sequence.
6. Transform: To transform a plasmid into a host, the CF line might look like:

   `Transform pDNA E.coli Amp

### Steps
PCR ca877F ca606R pAC-TetInv InvasinPCR
#Sequence information
ca877F GAGTTGGATCCNNNGGAGNNNNNNRTGNTGGTTTTCCAGCCAATCAGTG
ca606R GTCGACGGCGCTATTCAGATCCTC
pAC-TetInv GgatcGCGGCCGCtccctatcagtgatagagattgacatccctatcagtgatagagatactgagcacatcagcaggacgcactgaccgcatcagcaggacgcactgaccgGGATCCGTTTGACGTATGACAGGTATGCTTTATTTCATTTAAATTATGATGGTTTTCCAGCCAATCAGTGAGTTTCTCTTGATAAGGAATGCGGGAATGTCTATGTATTTTAATAAAATAATTTCATTTAATATTATTTCACGAATAGTTATTTGTATCTTTTTGATATGTGGAATGTTCATGGCTGGGGCTTCAGAAAAATATGATGCTAACGCACCGCAACAGGTCCAGCCTTATTCTGTCTCTTCATCTGCATTTGAAAATCTCCATCCTAATAATGAAATGGAGAGTTCAATCAATCCCTTTTCCGCATCGGATACAGAAAGAAATGCTGCAATAATAGATCGCGCCAATAAGGAGCAGGAGACTGAAGCGGTGAATAAGATGATAAGCACCGGGGCCAGGTTAGCTGCATCAGGCAGGGCATCTGATGTTGCTCACTCAATGGTGGGCGATGCGGTTAATCAAGAAATCAAACAGTGGTTAAATCGATTCGGTACGGCTCAAGTTAATCTGAATTTTGACAAAAATTTTTCGCTAAAAGAAAGCTCTCTTGATTGGCTGGCTCCTTGGTATGACTCTGCTTCATTCCTCTTTTTTAGTCAGTTAGGTATTCGCAATAAAGACAGCCGCAACACACTTAACCTTGGCGTCGGGATACGTACATTGGAGAACGGTTGGCTGTACGGACTTAATACTTTTTATGATAATGATTTGACCGGCCACAACCACCGTATCGGTCTTGGTGCCGAGGCCTGGACCGATTATTTACAGTTGGCTGCCAATGGGTATTTTCGCCTCAATGGATGGCACTCGTCGCGTGATTTCTCCGACTATAAAGAGCGCCCAGCCACTGGGGGGGATTTGCGCGCGAATGCTTATTTACCTGCACTCCCACAACTGGGGGGGAAGTTGATGTATGAGCAATACACCGGTGAGCGTGTTGCTTTATTTGGTAAAGATAATCTGCAACGCAACCCTTATGCCGTGACTGCCGGGATCAATTACACCCCCGTGCCTCTACTCACTGTCGGGGTAGATCAGCGTATGGGGAAAAGCAGTAAGCATGAAACACAGTGGAACCTCCAAATGAACTATCGCCTGGGCGAGAGTTTTCAGTCGCAACTTAGCCCTTCAGCGGTGGCAGGAACACGTCTACTGGCGGAGAGCCGCTATAACCTTGTCGATCGTAACAATAATATCGTGTTGGAGTATCAGAAACAGCAGGTGGTTAAACTGACATTATCGCCAGCAACTATCTCCGGCCTGCCGGGTCAGGTTTATCAGGTGAACGCACAAGTACAAGGGGCATCTGCTGTAAGGGAAATTGTCTGGAGTGATGCCGAACTGATTGCCGCTGGCGGCACATTAACACCACTGAGTACCACACAATTCAACTTGGTTTTACCGCCTTATAAACGCACAGCACAAGTGAGTCGGGTAACGGACGACCTGACAGCCAACTTTTATTCGCTTAGTGCGCTCGCGGTTGATCACCAAGGAAACCGATCTAACTCATTCACATTGAGCGTCACCGTTCAGCAGCCTCAGTTGACATTAACGGCGGCCGTCATTGGTGATGGCGCACCGGCTAATGGGAAAACTGCAATCACCGTTGAGTTCACCGTTGCTGATTTTGAGGGGAAACCCTTAGCCGGGCAGGAGGTGGTGATAACCACCAATAATGGTGCGCTACCGAATAAAATCACGGAAAAGACAGATGCAAATGGCGTCGCGCGCATTGCATTAACCAATACGACAGATGGCGTGACGGTAGTCACAGCAGAAGTGGAGGGGCAACGGCAAAGTGTTGATACCCACTTTGTTAAGGGTACTATCGCGGCGGATAAATCCACTCTGGCTGCGGTACCGACATCTATCATCGCTGATGGTCTAATGGCTTCAACCATCACGTTGGAGTTGAAGGATACCTATGGGGACCCGCAGGCTGGCGCGAATGTGGCTTTTGACACAACCTTAGGCAATATGGGCGTTATCACGGATCACAATGACGGCACTTATAGCGCACCATTGACCAGTACCACGTTGGGGGTAGCAACAGTAACGGTGAAAGTGGATGGGGCTGCGTTCAGTGTGCCGAGTGTGACGGTTAATTTCACGGCAGATCCTATTCCAGATGCTGGCCGCTCCAGTTTCACCGTCTCCACACCGGATATCTTGGCTGATGGCACGATGAGTTCCACATTATCCTTTGTCCCTGTCGATAAGAATGGCCATTTTATCAGTGGGATGCAGGGCTTGAGTTTTACTCAAAACGGTGTGCCGGTGAGTATTAGCCCCATTACCGAGCAGCCAGATAGCTATACCGCGACGGTGGTTGGGAATAGTGTCGGTGATGTCACAATCACGCCGCAGGTTGATACCCTGATACTGAGTACATTGCAGAAAAAAATATCCCTATTCCCGGTACCTACGCTGACCGGTATTCTGGTTAACGGGCAAAATTTCGCTACGGATAAAGGGTTCCCGAAAACGATCTTTAAAAACGCCACATTCCAGTTACAGATGGATAACGATGTTGCTAATAATACTCAGTATGAGTGGTCGTCGTCATTCACACCCAATGTATCGGTTAACGATCAGGGTCAGGTGACGATTACCTACCAAACCTATAGCGAAGTGGCTGTGACGGCGAAAAGTAAAAAATTCCCAAGTTATTCGGTGAGTTATCGGTTCTACCCAAATCGGTGGATATACGATGGCGGCAGATCGCTGGTATCCAGTCTCGAGGCCAGCAGACAATGCCAAGGTTCAGATATGTCTGCGGTTCTTGAATCCTCACGTGCAACCAACGGAACGCGTGCGCCTGACGGGACATTGTGGGGCGAGTGGGGGAGCTTGACCGCGTATAGTTCTGATTGGCAATCTGGTGAATATTGGGTCAAAAAGACCAGCACGGATTTTGAAACCATGAATATGGACACAGGCGCACTGCAACCAGGGCCTGCATACTTGGCGTTCCCGCTCTGTGCGCTGTCAATATAAgaattcgaagcttgggcccgaacaaaaactcatctcagaagaggatctgaatagcgccgtcgaccatcatcatcatcatcattgagtttaaacggtctccagcttggctgttttggcggatgagagaagattttcagcctgatacagattaaatcagaacgcagaagcggtctgataaaacagaatttgcctggcggcagtagcgcggtggtcccacctgaccccatgccgaactcagaagtgaaacgccgtagcgccgatggtagtgtggggtctccccatgcgagagtagggaactgccaggcatcaaataaaacgaaaggctcagtcgaaagactgggcctttcgttttatctgttgtttgtcggtgaactaattcgaagcttgacataagcggctatttaacgaccctgccctgaaccgacgaccgggtcgaatttgctttcgaatttctgccattcatccgcttattatcacttattcaggcgtagcaccaggcgtttaagggcaccaataactgccttaaaaaaattacgccccgccctgccactcatcgcagtactgttgtaattcattaagcattctgccgacatggaagccatcacagacggcatgatgaacctgaatcgccagcggcatcagcaccttgtcgccttgcgtataatatttgcccatcgtgaaaacgggggcgaagaagttgtccatattggccacgtttaaatcaaaactggtgaaactcacccagggattggctgagacgaaaaacatattctcaataaaccctttagggaaataggccaggttttcaccgtaacacgccacatcttgcgaatatatgtgtagaaactgccggaaatcgtcgtggtattcactccagagcgatgaaaacgtttcagtttgctcatggaaaacggtgtaacaagggtgaacactatcccatatcaccagctcaccgtctttcattgccatacggaactccggatgagcattcatcaggcgggcaagaatgtgaataaaggccggataaaacttgtgcttatttttctttacggtctttaaaaaggccgtaatatccagctgaacggtctggttataggtacattgagcaactgactgaaatgcctcaaaatgttctttacgatgccattgggatatatcaacggtggtatatccagtgatttttttctccattttagcttccttagctcctgaaaatctcgataactcaaaaaatacgcccggtagtgatcttatttcattatggtgaaagttggaacctcttacgtgccgatcaacgtctcattttcgccaaaagttggcccagggcttcccggtatcaacagggacaccaggatttatttattctgcgaagtgatcttccgtcacaggtatttattcggcgcaaagtgcgtcgggtgatgctgccaacttactgatttagtgtatgatggtgtttttgaggtgctccagtggcttctgtttctatcagctgtccctcctgttcagctactgacggggtggtgcgtaacggcaaaagcaccgccggacatcagcgctagcggagtgtatactggcttactatgttggcactgatgagggtgtcagtgaagtgcttcatgtggcaggagaaaaaaggctgcaccggtgcgtcagcagaatatgtgatacaggatatattccgcttcctcgctcactgactcgctacgctcggtcgttcgactgcggcgagcggaaatggcttacgaacggggcggagatttcctggaagatgccaggaagatacttaacagggaagtgagagggccgcggcaaagccgtttttccataggctccgcccccctgacaagcatcacgaaatctgacgctcaaatcagtggtggcgaaacccgacaggactataaagataccaggcgtttccccctggcggctccctcgtgcgctctcctgttcctgcctttcggtttaccggtgtcattccgctgttatggccgcgtttgtctcattccacgcctgacactcagttccgggtaggcagttcgctccaagctggactgtatgcacgaaccccccgttcagtccgaccgctgcgccttatccggtaactatcgtcttgagtccaacccggaaagacatgcaaaagcaccactggcagcagccactggtaattgatttagaggagttagtcttgaagtcatgcgccggttaaggctaaactgaaaggacaagttttggtgactgcgctcctccaagccagttacctcggttcaaagagttggtagctcagagaaccttcgaaaaaccgccctgcaaggcggttttttcgttttcagagcaagagattacgcgcagaccaaaacgatctcaagaagatcatcttattaatcagataaaatatttctagatttcagtgcaatttatctcttcaaatgtagcacctgaagtc
What is the sequence of the pcr product?

PCR, or Polymerase Chain Reaction, is a method used to amplify a specific region of DNA. It uses a pair of primers that flank the target region to produce copies of that region.

The sequence of the PCR product depends on the sequences of the primers and the template DNA.

The primers in this case are ca877F and ca606R, and the template DNA is pAC-TetInv.

The ca877F primer sequence is:
GAGTTGGATCCNNNGGAGNNNNNNRTGNTGGTTTTCCAGCCAATCAGTG

The ca606R primer sequence is:
GTCGACGGCGCTATTCAGATCCTC

The primer sequences are used to start the replication process. The forward primer (ca877F) binds to the start of the sequence to be amplified, and the reverse primer (ca606R) binds to the end of the sequence to be amplified. The template is read from 3' to 5', and new DNA is synthesized from 5' to 3'.

To find the sequence of the PCR product, you have to find the region on the template DNA that is flanked by the primer sequences. However, your provided primers contain ambiguous bases (represented by N and R), which makes it impossible to accurately identify the exact location of the primers on the template sequence.

If the locations of the primers could be determined, the PCR product would be the sequence on the template DNA that lies between the locations of the primers, including the primer sequences themselves.

Due to the complexity of this task, it may be best to use a bioinformatics software or online tool to accurately find the sequence of the PCR product.

Please note that the PCR process doesn't change the sequence of the amplified DNA. It only produces more copies of the same sequence. Therefore, the sequence of the PCR product will be identical to the sequence of the target region on the template DNA.

Why can't you predict the product directly?

Determining the exact sequence of a PCR product requires a few more pieces of information than what has been provided. Let's break down what we need:

1. **Template sequence:** The exact sequence of the DNA template you are using is necessary. The primers bind to this template sequence to initiate the PCR.
2. **Primer sequences:** The ca877F and ca606R sequences have been provided. These bind to the DNA template and determine the endpoints of the PCR product.
3. **Strand orientation:** Knowing which strand (5'-3' or 3'-5') the primers bind to is also important. Primers always bind in the 3'-5' direction on the template strand, which allows new DNA to be synthesized in the 5'-3' direction.
4. **Primer locations:** It's necessary to know where exactly these primers bind on your DNA template. If they are not part of the template sequence provided, then it becomes difficult to predict the PCR product.

It seems that the ca877F and ca606R primers do not match the provided sequence of pAC-TetInv. Therefore, it's not possible to predict the PCR product from the information given. If you have the template sequence that these primers are designed to bind, and know their binding locations, then it would be possible to determine the sequence of the PCR product.
