## Supplementary material for "Enabling AI in Synthetic Biology through Construction File Specification": All supplemental files: pydna_plugin_test.html

PyDNA\_CF\_Simulator plugin test - A ShareGPT conversation 

Model: Plugins
•
Enabled plugins:

simulate this exactly as is:
### Define sequences
forward CCGCAACACACTTAACCTTG
reverse GTGGTTGTGGCCGGTCAAATC
template CCGCAACACACTTAACCTTGGCGTCGGGATACGTACATTGGAGAACGGTTGGCTGTACGGACTTAATACTTTTTATGATAATGATTTGACCGGCCACAACCACCG
### Define steps
PCR forward reverse template product

The simulation of your PCR process was successful. Here are the details of the sequences involved:

1. Forward Primer:

   - Sequence: CCGCAACACACTTAACCTTG
   - 5' Modification: Hydroxyl
   - 3' Modification: None
   - Is Circular: No
   - Is Double Stranded: No
2. Reverse Primer:

   - Sequence: GTGGTTGTGGCCGGTCAAATC
   - 5' Modification: Hydroxyl
   - 3' Modification: None
   - Is Circular: No
   - Is Double Stranded: No
3. Template:

   - Sequence: CCGCAACACACTTAACCTTGGCGTCGGGATACGTACATTGGAGAACGGTTGGCTGTACGGACTTAATACTTTTTATGATAATGATTTGACCGGCCACAACCACCG
   - 5' Modification: None
   - 3' Modification: None
   - Is Circular: Yes
   - Is Double Stranded: Yes
4. Product:

   - Sequence: CCGCAACACACTTAACCTTGGCGTCGGGATACGTACATTGGAGAACGGTTGGCTGTACGGACTTAATACTTTTTATGATAATGATTTGACCGGCCACAACCAC
   - 5' Modification: Hydroxyl
   - 3' Modification: Hydroxyl
   - Is Circular: No
   - Is Double Stranded: Yes

Please let me know if you need further assistance.
