## Supplementary material for "Enabling AI in Synthetic Biology through Construction File Specification": All supplemental files: syntax_conversions.html

CF Syntax Conversions - A ShareGPT conversation 

Model: GPT-4

### CF Shorthand Specification
#### Layer 0: Basic Concept
A Construction File (CF) is a list of Steps, with each Step representing a specific operation in a molecular biology experiment. Each Step is written on a new line. Parameters are separate by whitespace, preferably TSV. The Step includes the names of input DNA sequence(s), non-sequence parameters, and ends with the name of the product DNA sequence. The input sequences can refer to products of previous steps. The product is the output of the operation.
In addition, a CF can include sequences in the form of 'name sequence' lines.
Comment lines follow '#', '//', or '/\*comment\*/' syntax.
#### Layer 1: Core Operations
This layer defines a core set of operations: PCR, Digest, Ligate, GoldenGate, Gibson, and Transform. Each operation has specific parameters and (optionally) a product name.
#### Examples
- `PCR ForwardPrimer ReversePrimer Template ProductName`
- ForwardPrimer, ReversePrimer, Template: Names of DNA sequences
- `Digest DNA Enzymes FragmentSelection ProductName`
- DNA: Name of the DNA sequence to be digested
- Enzymes: List of enzymes to be used, e.g. EcoRI,BamHI
- FragmentSelection: Index indicating the chosen fragment post-digestion, according to orientation and origin given in the input DNA.
- `Ligate Fragment1 Fragment2 ProductName`
- Fragment1, Fragment2: Names of DNA sequences
- `GoldenGate Fragment1 Fragment2 Enzyme ProductName`
- Fragment1, Fragment2: Names of DNA sequences
- Enzyme: Type IIS-like enzyme to be used along with ligase. e.g. BsaI
- `Gibson Fragment1 Fragment2 ProductName`
- Fragment1, Fragment2: Names of DNA sequences
- `Transform Plasmid Host Antibiotic Temperature ProductName`
- Plasmid: Name of the DNA sequence
- Host: Bacterial strain
- Antibiotics: Antibiotics used, e.g. Amp,Kan
- Temperature: Incubation temperature in Celsius, e.g. 37
Here is an example I wish to discuss:
//Figure 1 Example: PCR, cut, and ligate with palindromic type II six-cutters of two fragments
//Derived from https://openwetware.org/wiki/Arking:JCAOligoTutorial1
### Cloning Steps
PCR ca1067F ca1067R pSB1AK3-b0015 pcrpdt
Digest pcrpdt EcoRI,SpeI 1 pcrdig
Digest pSB1A2-I13521 EcoRI,SpeI 1 vectdig
Ligate pcrdig vectdig lig
Transform lig DH10B Amp 37 pSB1A2-Bca9128

This is a simplified way to describe the steps of a molecular cloning experiment in a format that is easy to understand and programmatically analyze. Each line represents a step in the process, and parameters are clearly separated by whitespace, making it easy to parse.

The process in your example involves five steps:

1. **PCR (Polymerase Chain Reaction):** This step involves the amplification of a specific DNA sequence using forward and reverse primers (ca1067F and ca1067R). The template DNA sequence for the PCR is pSB1AK3-b0015, and the product (output) of this reaction is named as pcrpdt.
2. **Digestion of PCR product:** The PCR product is then digested by the restriction enzymes EcoRI and SpeI. In this case, the first fragment that appears post-digestion (fragment selection=1) is chosen, and the product is named pcrdig.
3. **Digestion of vector:** A vector sequence (pSB1A2-I13521) is also digested by the same enzymes, EcoRI and SpeI. Again, the first fragment post-digestion is chosen, and the resulting product is named vectdig.
4. **Ligation:** The digested PCR product (pcrdig) and the digested vector (vectdig) are then ligated together in this step. The product of this ligation reaction is named lig.
5. **Transformation:** The ligated DNA (lig) is then transformed into a strain of bacteria called DH10B, using ampicillin (Amp) as the antibiotic and incubating at a temperature of 37 degrees Celsius. The resulting product, presumably a colony of bacteria containing the plasmid, is named pSB1A2-Bca9128.

It is important to note that this is a very simplified representation of the process, and each step would actually involve a number of sub-steps and more specific parameters in a real experimental setup. The format, however, is useful for planning and documenting molecular biology experiments, and for providing a clear overview of the cloning process.

Express it according to this json spec:
/\*\*
\* A Construction File (CF) is a structured format for specifying a series of molecular biology operations.
\* CFs are used to plan, simulate, and document experiments.
\*
\* Syntax:
\* A Construction File is represented as a JSON object, containing two main elements: 'steps'
\* and 'sequences'. The 'steps' is an array of objects, where each object represents a construction
\* step with its associated operation, input sequences, and output product. The 'sequences' is an object
\* containing key-value pairs, where each key is a unique identifier for a DNA sequence, and the value is the
\* actual sequence as a string.
\*
\* Example:
\* Here's a simple example of a Construction File that demonstrates PCR and GoldenGate steps.
\*
\* {
\* "steps": [
\* {
\* "operation": "PCR",
\* "output": "P6",
\* "forward\_oligo": "P6libF",
\* "reverse\_oligo": "P6libR",
\* "template": "pTP1",
\* "product\_size": 3583
\* },
\* {
\* "operation": "GoldenGate",
\* "dnas": ["P6"],
\* "enzyme": "BsaI",
\* "output": "pP6"
\* }
\* ],
\* "sequences": {
\* "P6libF": {"sequence": "ccaaaggtctcATTATANNNNNNNNNNNNNNNNNTGTCAANNNNGAacccaggactcctcgaagtcgttcttaagacaac", "ext5": null, "ext3": null, "is\_double\_stranded": false, "is\_circular": false, "mod\_ext5": "hydroxyl", "mod\_ext3": null},
\* "P6libR": {"sequence": "cagttGGTCTCAATAATNNNNNNANNNNGTtagtatttctcctcgtctacggttaactgatactc", "ext5": null, "ext3": null, "is\_double\_stranded": false, "is\_circular": false, "mod\_ext5": "hydroxyl", "mod\_ext3": null},
\* "pTP1": {"sequence": "ATTACCGCCTTTGAGTGG", "ext5": "", "ext3": "", "is\_double\_stranded": true, "is\_circular": true, "mod\_ext5": null, "mod\_ext3": null}
\* }
\* }
\*
\* In this example, the 'steps' array has two steps: PCR and Assemble. The PCR step uses forward and
\* reverse oligos "P6libF" and "P6libR", with plasmid "pTP1" as the template. The PCR product is named "P6". The Assemble
\* step uses the "P6" PCR product and "BsaI"-based Golden Gate Assembly to create a final output named "pP6". The 'sequences'
\* object contains the sequences for "P6libF", "P6libR", and "pTP1" as Polynucleotide objects.
\*
\* @typedef {Object} ConstructionFile
\* @property {Array.} steps - An array of construction steps, where each step is an operation object.
\* @property {Object.} sequences - An object containing key-value pairs of sequence names and their corresponding Polynucleotide objects.
\*
\* @typedef {Object} Polynucleotide
\* @property {string} sequence - The DNA sequence following the regex pattern /^[ATCGNRKYSWBVHDM]+$/.
\* @property {string|null} ext5 - The 'left' end's extension, which can be null or a string following the regex pattern /^(-)?[ATCG]+$/.
\* @property {string|null} ext3 - The 'right' end's extension, which can be null or a string following the regex pattern /^(-)?[ATCG]+$/.
\* @property {boolean} is\_double\_stranded - Indicates if the polynucleotide is double-stranded.
\* @property {boolean} is\_circular - Indicates if the polynucleotide is circular.
\* @property {string|null} mod\_ext5 - The 5' modification of coding strand.
\* @property {string|null} mod\_ext3 - The 5' modification of non-coding strand.
\*
\* @typedef {Object} PCR
\* @property {'PCR'} operation - The type of operation.
\* @property {string} forward\_oligo - The name of the forward primer used in PCR operation (DNA identifier).
\* @property {string} reverse\_oligo - The name of the reverse primer used in PCR operation (DNA identifier).
\* @property {string} template - The name of the template DNA used in PCR operation (DNA identifier).
\* @property {string} output - The name of the output product of the operation (DNA identifier).
\* @property {number|undefined} [product\_size] - The expected product size in PCR operation (optional).
\*
\* @typedef {Object} GoldenGate
\* @property {'GoldenGate'} operation - The type of operation.
\* @property {Array.} dnas - An array of DNA part identifiers being assembled in the GoldenGate reaction.
\* @property {'AarI'|'BbsI'|'BsaI'|'BsmBI'|'SapI'|'BseRI'} enzyme - The Type IIS-like enzyme used with ligase in the assembly reaction.
\* @property {string} output - The name of the output product of the operation (DNA identifier).
\*
\* @typedef {Object} Gibson
\* @property {'Gibson'} operation - The type of operation.
\* @property {Array.} dnas - An array of DNA part identifiers being assembled using the Gibson reaction.
\* @property {string} output - The name of the output product of the operation (DNA identifier).
\*
\* @typedef {Object} Transform
\* @property {'Transform'} operation - The type of operation.
\* @property {string} dna - The identifier of the DNA being introduced into cells.
\* @property {string} strain - The bacterial strain being transformed with the DNA.
\* @property {'Amp'|'Carb'|'Cam'|'Kan'|'Gen'|'Spec'|'Trim'} antibiotics - The antibiotics used in the growth medium.
\* @property {number|undefined} [temperature] - The temperature used to grow the transformed cells (optional).
\* @property {string} output - The name of the output cytoplasmic DNAs (DNA identifier).
\*
\* @typedef {Object} Digest
\* @property {'Digest'} operation - The type of operation.
\* @property {string} dna - The identifier of the DNA used in the Digest operation.
\* @property {(Array.&lt;'AarI'|'BbsI'|'BsaI'|'BsmBI'|'SapI'|'BseRI'|'BamHI'|'BglII'|'EcoRI'|'XhoI'|'SpeI'|'XbaI'|'PstI'|'HindIII'|'NotI'|'XmaI'|'SmaI'|'KpnI'|'SacI'|'Sal'&gt;|Array.)} enzymes - The enzymes used in the Digest operation.
\* @property {number} fragSelect - The index, counted from zero, of the output fragment.
\* @property {string} output - The name of the output product of the operation (DNA identifier).
\* @property {number|undefined} [product\_size] - The expected size of the selected fragment (optional).
\*
\* @typedef {Object} Ligate
\* @property {'Ligate'} operation - The type of operation.
\* @property {Array.} dnas - An array of DNA part identifiers for digested DNAs being joined in the Ligate reaction.
\* @property {string} output - The name of the output product of the operation (DNA identifier).
\*/

Sure, the Construction File for the specified steps in JSON format would look something like this:

```
jsonCopy code


{
    "steps": [
        {
            "operation": "PCR",
            "forward_oligo": "ca1067F",
            "reverse_oligo": "ca1067R",
            "template": "pSB1AK3-b0015",
            "output": "pcrpdt"
        },
        {
            "operation": "Digest",
            "dna": "pcrpdt",
            "enzymes": ["EcoRI","SpeI"],
            "fragSelect": 1,
            "output": "pcrdig"
        },
        {
            "operation": "Digest",
            "dna": "pSB1A2-I13521",
            "enzymes": ["EcoRI","SpeI"],
            "fragSelect": 1,
            "output": "vectdig"
        },
        {
            "operation": "Ligate",
            "dnas": ["pcrdig", "vectdig"],
            "output": "lig"
        },
        {
            "operation": "Transform",
            "dna": "lig",
            "strain": "DH10B",
            "antibiotics": "Amp",
            "temperature": 37,
            "output": "pSB1A2-Bca9128"
        }
    ],
    "sequences": {
        "ca1067F": {
            "sequence": "<sequence of ca1067F>",
            "ext5": null,
            "ext3": null,
            "is_double_stranded": false,
            "is_circular": false,
            "mod_ext5": "hydroxyl",
            "mod_ext3": null
        },
        "ca1067R": {
            "sequence": "<sequence of ca1067R>",
            "ext5": null,
            "ext3": null,
            "is_double_stranded": false,
            "is_circular": false,
            "mod_ext5": "hydroxyl",
            "mod_ext3": null
        },
        "pSB1AK3-b0015": {
            "sequence": "<sequence of pSB1AK3-b0015>",
            "ext5": null,
            "ext3": null,
            "is_double_stranded": true,
            "is_circular": true,
            "mod_ext5": null,
            "mod_ext3": null
        },
        "pSB1A2-I13521": {
            "sequence": "<sequence of pSB1A2-I13521>",
            "ext5": null,
            "ext3": null,
            "is_double_stranded": true,
            "is_circular": true,
            "mod_ext5": null,
            "mod_ext3": null
        }
    }
}
```

Please note that I have placed placeholder `<sequence of ...>` where the actual DNA sequences would be because I do not have those sequences. Please replace them with the actual DNA sequences. The value of `is_double_stranded` and `is_circular` will vary depending on the actual sequences. Similarly, the 5' modifications (`mod_ext5` and `mod_ext3`) are placeholders and should be replaced with the correct modifications if any exist.

Express it as XML

Certainly, here's the equivalent representation of the steps in XML format:

```
xmlCopy code


<ConstructionFile>
    <Steps>
        <Step>
            <Operation>PCR</Operation>
            <ForwardOligo>ca1067F</ForwardOligo>
            <ReverseOligo>ca1067R</ReverseOligo>
            <Template>pSB1AK3-b0015</Template>
            <Output>pcrpdt</Output>
        </Step>
        <Step>
            <Operation>Digest</Operation>
            <DNA>pcrpdt</DNA>
            <Enzymes>EcoRI,SpeI</Enzymes>
            <FragSelect>1</FragSelect>
            <Output>pcrdig</Output>
        </Step>
        <Step>
            <Operation>Digest</Operation>
            <DNA>pSB1A2-I13521</DNA>
            <Enzymes>EcoRI,SpeI</Enzymes>
            <FragSelect>1</FragSelect>
            <Output>vectdig</Output>
        </Step>
        <Step>
            <Operation>Ligate</Operation>
            <DNAs>
                <DNA>pcrdig</DNA>
                <DNA>vectdig</DNA>
            </DNAs>
            <Output>lig</Output>
        </Step>
        <Step>
            <Operation>Transform</Operation>
            <DNA>lig</DNA>
            <Strain>DH10B</Strain>
            <Antibiotics>Amp</Antibiotics>
            <Temperature>37</Temperature>
            <Output>pSB1A2-Bca9128</Output>
        </Step>
    </Steps>
    <Sequences>
        <Sequence id="ca1067F">
            <SequenceData><sequence of ca1067F></SequenceData>
            <Ext5></Ext5>
            <Ext3></Ext3>
            <IsDoubleStranded>false</IsDoubleStranded>
            <IsCircular>false</IsCircular>
            <ModExt5>hydroxyl</ModExt5>
            <ModExt3></ModExt3>
        </Sequence>
        <Sequence id="ca1067R">
            <SequenceData><sequence of ca1067R></SequenceData>
            <Ext5></Ext5>
            <Ext3></Ext3>
            <IsDoubleStranded>false</IsDoubleStranded>
            <IsCircular>false</IsCircular>
            <ModExt5>hydroxyl</ModExt5>
            <ModExt3></ModExt3>
        </Sequence>
        <Sequence id="pSB1AK3-b0015">
            <SequenceData><sequence of pSB1AK3-b0015></SequenceData>
            <Ext5></Ext5>
            <Ext3></Ext3>
            <IsDoubleStranded>true</IsDoubleStranded>
            <IsCircular>true</IsCircular>
            <ModExt5></ModExt5>
            <ModExt3></ModExt3>
        </Sequence>
        <Sequence id="pSB1A2-I13521">
            <SequenceData><sequence of pSB1A2-I13521></SequenceData>
            <Ext5></Ext5>
            <Ext3></Ext3>
            <IsDoubleStranded>true</IsDoubleStranded>
            <IsCircular>true</IsCircular>
            <ModExt5></ModExt5>
            <ModExt3></ModExt3>
        </Sequence>
    </Sequences>
</ConstructionFile>
```

As in the previous JSON response, I have placed placeholder `<sequence of ...>` where the actual DNA sequences would be because I do not have those sequences. Please replace them with the actual DNA sequences. The value of `IsDoubleStranded` and `IsCircular` will vary depending on the actual sequences. Similarly, the 5' modifications (`ModExt5` and `ModExt3`) are placeholders and should be replaced with the correct modifications if any exist.
