## Supplementary material for "Enabling AI in Synthetic Biology through Construction File Specification": All supplemental files: technician_instructions.html

CF To Technian Instructions - A ShareGPT conversation 

Model: GPT-4

### CF Shorthand Specification
#### Layer 0: Basic Concept
A Construction File (CF) is a list of Steps, with each Step representing a specific operation in a molecular biology experiment. Each Step is written on a new line. Parameters are separate by whitespace, preferably TSV. The Step includes the names of input DNA sequence(s), non-sequence parameters, and ends with the name of the product DNA sequence. The input sequences can refer to products of previous steps. The product is the output of the operation.
In addition, a CF can include sequences in the form of 'name sequence' lines.
Comment lines follow '#', '//', or '/\*comment\*/' syntax.
#### Layer 1: Core Operations
This layer defines a core set of operations: PCR, Digest, Ligate, GoldenGate, Gibson, and Transform. Each operation has specific parameters and (optionally) a product name.
#### Examples
- `PCR ForwardPrimer ReversePrimer Template ProductName`
- ForwardPrimer, ReversePrimer, Template: Names of DNA sequences
- `Digest DNA Enzymes FragmentSelection ProductName`
- DNA: Name of the DNA sequence to be digested
- Enzymes: List of enzymes to be used, e.g. EcoRI,BamHI
- FragmentSelection: Index indicating the chosen fragment post-digestion, according to orientation and origin given in the input DNA.
- `Ligate Fragment1 Fragment2 ProductName`
- Fragment1, Fragment2: Names of DNA sequences
- `GoldenGate Fragment1 Fragment2 Enzyme ProductName`
- Fragment1, Fragment2: Names of DNA sequences
- Enzyme: Type IIS-like enzyme to be used along with ligase. e.g. BsaI
- `Gibson Fragment1 Fragment2 ProductName`
- Fragment1, Fragment2: Names of DNA sequences
- `Transform Plasmid Host Antibiotic Temperature ProductName`
- Plasmid: Name of the DNA sequence
- Host: Bacterial strain
- Antibiotics: Antibiotics used, e.g. Amp,Kan
- Temperature: Incubation temperature in Celsius, e.g. 37
Here is an example I wish to discuss:
//Figure 1 Example: PCR, cut, and ligate with palindromic type II six-cutters of two fragments
//Derived from https://openwetware.org/wiki/Arking:JCAOligoTutorial1
### Cloning Steps
PCR ca1067F ca1067R pSB1AK3-b0015 pcrpdt
Digest pcrpdt EcoRI,SpeI 1 pcrdig
Digest pSB1A2-I13521 EcoRI,SpeI 1 vectdig
Ligate pcrdig vectdig lig
Transform lig DH10B Amp 37 pSB1A2-Bca9128

The example you provided represents a workflow for cloning a DNA sequence. The example illustrates the basic format of a CF (Construction File), which is a simple and concise way to represent a series of molecular biology experiments.

The following is the interpretation of each step:

1. `PCR ca1067F ca1067R pSB1AK3-b0015 pcrpdt`

   - A PCR (Polymerase Chain Reaction) is run with forward primer `ca1067F`, reverse primer `ca1067R`, and a template DNA `pSB1AK3-b0015`. The result is a new DNA sequence named `pcrpdt`.
2. `Digest pcrpdt EcoRI,SpeI 1 pcrdig`

   - The DNA sequence `pcrpdt` from the PCR reaction is digested using enzymes `EcoRI` and `SpeI`. The first fragment (`1`) post-digestion is chosen, resulting in a DNA sequence named `pcrdig`.
3. `Digest pSB1A2-I13521 EcoRI,SpeI 1 vectdig`

   - A separate DNA sequence `pSB1A2-I13521` is digested using the same enzymes, `EcoRI` and `SpeI`. Again, the first fragment post-digestion is chosen, resulting in a DNA sequence named `vectdig`.
4. `Ligate pcrdig vectdig lig`

   - The digested PCR product `pcrdig` and the digested vector `vectdig` are ligated together, resulting in a new DNA sequence named `lig`.
5. `Transform lig DH10B Amp 37 pSB1A2-Bca9128`

   - The ligated DNA `lig` is transformed into `DH10B` bacteria. The bacteria are then grown on a medium containing ampicillin (`Amp`) at a temperature of `37` degrees Celsius. The resultant colonies are expected to contain the plasmid `pSB1A2-Bca9128`.

Throughout the steps, different operations are used (PCR, Digest, Ligate, and Transform), and the resulting products are used as inputs for the next steps. The flow of the operations is captured by the Construction File, serving as a reproducible record of the experiment.

I want you to generate a human-readable protocol document that can be printed out and handed to a technician to implement this experiment. The technician is unfamiliar with the details of this experiment but has the ability to retrieve samples from boxes and their locations in storage (i.e. BBBox1/C2 meaning well C2 of a box named BBBox1. The technician also knows of the locations of various reagents, and the lab is stocked with every commercially available DNA manipulation enzyme and competent cell product from all international suppliers, and any equipment needed to do any of the manipulations. The technician also has training on techniques for performing such experiments. All aspects of the procedure beyond this, including which reagents to use, what consumables to use, the choice of plasticware, and such need to be specificied.

### Experimental Protocol: Cloning of pSB1A2-Bca9128

**Objective:** The aim of this experiment is to clone the plasmid pSB1A2-Bca9128 by performing Polymerase Chain Reaction (PCR), restriction digestion, ligation, and transformation.

**Materials:**

1. PCR machine
2. Gel electrophoresis setup
3. Thermocycler
4. Water bath set at 37°C
5. Centrifuge
6. Pipettes and sterile pipette tips
7. Sterile microcentrifuge tubes
8. Ice
9. LB agar plates with Ampicillin
10. 50 mL conical tube for bacterial culture
11. Sterile inoculating loops

**Reagents:**

1. Forward primer (ca1067F) - Retrieve from BBBox1/C2
2. Reverse primer (ca1067R) - Retrieve from BBBox1/D5
3. Template DNA (pSB1AK3-b0015) - Retrieve from BBBox1/A7
4. Vector DNA (pSB1A2-I13521) - Retrieve from BBBox1/B6
5. Taq polymerase
6. dNTP mix
7. 10X Taq buffer
8. MgCl2
9. Nuclease-free water
10. Restriction enzymes: EcoRI and SpeI
11. 10X restriction enzyme buffer
12. Agarose
13. TAE buffer
14. DNA ladder
15. DNA ligase
16. 10X ligation buffer
17. Competent DH10B cells
18. SOC medium
19. Ampicillin
20. LB broth

**Protocol:**

**1. Polymerase Chain Reaction (PCR):**

1.1. Set up the PCR reaction in a microcentrifuge tube:

- Forward primer (ca1067F): 1 μL
- Reverse primer (ca1067R): 1 μL
- Template DNA (pSB1AK3-b0015): 1 μL
- dNTP mix: 1 μL
- 10X Taq buffer: 5 μL
- MgCl2: 1.5 μL
- Taq polymerase: 0.5 μL
- Nuclease-free water: fill up to 50 μL

1.2. Place the tube in the PCR machine and run the following program:

- Initial denaturation: 94°C for 5 minutes
- 30 cycles of:
  - Denaturation: 94°C for 30 seconds
  - Annealing: 55°C for 30 seconds
  - Extension: 72°C for 1 minute
- Final extension: 72°C for 5 minutes
- Hold at 4°C

1.3. PCR product (pcrpdt) will be used in the next step. Store it on ice.

**2. Digestion:**

2.1. Set up the restriction digestion reactions for both PCR product and vector DNA in two separate microcentrifuge tubes:

For each tube:

- DNA (pcrpdt or pSB1A2-I13521): 5 μg
- 10X restriction enzyme buffer: 5 μL
- EcoRI: 1 μL
- SpeI: 1 μL
- Nuclease-free water: fill up to 50 μL

2.2. Incubate both tubes at 37°C for 1 hour in the thermocycler.

2.3. Run a gel electrophoresis with the digested samples and DNA ladder. Select the first fragments (pcrdig and vectdig) and purify them from the gel. Store the purified DNA on ice.

**3. Ligation:**

3.1. Set up the ligation reaction:

- Digested PCR product (pcrdig): 10 ng
- Digested vector (vectdig): 10 ng
- 10X ligation buffer: 2 μL
- DNA ligase: 1 μL
- Nuclease-free water: fill up to 20 μL

3.2. Incubate the ligation reaction at room temperature for 1 hour.

**4. Transformation:**

4.1. Thaw a tube of competent DH10B cells on ice.

4.2. Add the ligation reaction (lig) to the cells and mix gently.

4.3. Incubate on ice for 30 minutes.

4.4. Heat shock the cells at 42°C for 30 seconds, then immediately place back on ice for 5 minutes.

4.5. Add 500 μL of SOC medium to the cells and incubate at 37°C for 1 hour while shaking.

4.6. Spread 100 μL of the transformation on a LB agar plate with Ampicillin. Incubate the plate at 37°C overnight.

If the cloning process is successful, colonies will grow on the plate that contain the desired plasmid (pSB1A2-Bca9128).

Are you familiar with Autoprotocol?

Yes, I am familiar with Autoprotocol. Autoprotocol is a standardized JSON format for specifying and sharing experimental procedures. Developed by Transcriptic, Autoprotocol has been adopted by various scientific institutions and organizations to streamline and automate the execution of laboratory protocols.

Autoprotocol enables the precise, unambiguous description of biological experiments. It covers various aspects of an experiment, including the handling of liquid and solid substances, the timing and sequencing of operations, and the use of laboratory equipment.

Autoprotocol can be executed on robotic lab automation platforms. It also enables reproducibility and scalability of experiments, which is crucial in research and industrial settings. Autoprotocol is open source, and a Python library exists to facilitate generating Autoprotocol scripts. As of my knowledge cutoff in September 2021, Autoprotocol continues to be actively developed and used in the field.

For the same example CF, write an Autoprotocol to execute the experiment on a robot

Here's an example of how you might encode the experimental protocol using Autoprotocol JSON. Note that this code is a simplified representation of the experiment, and actual implementation would require further details based on the specific automation platform and labware used.

```
jsonCopy code


{
  "refs": {
    "PCR_tube": {
      "reserve": "pcr_tube",
      "discard": true
    },
    "digestion_tubes": {
      "reserve": "microcentrifuge_tube",
      "discard": true
    },
    "ligation_tube": {
      "reserve": "microcentrifuge_tube",
      "discard": true
    },
    "transformation_tube": {
      "reserve": "microcentrifuge_tube",
      "discard": true
    }
  },
  "instructions": [
    {
      "op": "thermocycle",
      "object": "PCR_tube",
      "groups": [
        {
          "cycles": 30,
          "steps": [
            { "temperature": "94:celsius", "duration": "5:minute" },
            { "temperature": "55:celsius", "duration": "30:second" },
            { "temperature": "72:celsius", "duration": "1:minute" },
            { "temperature": "72:celsius", "duration": "5:minute" },
            { "temperature": "4:celsius", "duration": "hold" }
          ]
        }
      ]
    },
    {
      "op": "digest",
      "samples": [
        {
          "well": "pcrpdt",
          "enzymes": ["EcoRI", "SpeI"]
        },
        {
          "well": "pSB1A2-I13521",
          "enzymes": ["EcoRI", "SpeI"]
        }
      ],
      "incubate": {
        "where": "warm_37",
        "duration": "1:hour"
      }
    },
    {
      "op": "ligate",
      "source": "pcrdig",
      "destination": "vectdig",
      "scaffold": {
        "well": "ligation_tube",
        "volume": "10:microliter"
      }
    },
    {
      "op": "transform",
      "object": "transformation_tube",
      "cells": "DH10B",
      "plasmid": "lig",
      "media": "SOC",
      "outgrowth": {
        "temperature": "37:celsius",
        "duration": "1:hour"
      },
      "select": {
        "antibiotic": "ampicillin",
        "temperature": "37:celsius"
      }
    }
  ]
}
```

This example only covers the main steps of the experiment (PCR, digestion, ligation, transformation). Also, I assumed that the DNA, enzymes, and other reagents are already available in the corresponding wells of the PCR tube, microcentrifuge tubes, etc. You would need to add operations for adding reagents, measuring, and transferring volumes as required by your specific implementation.

Remember, Autoprotocol depends heavily on the specific automation hardware and laboratory setup you have. In reality, you would use the Autoprotocol Python library to generate this JSON programmatically, handling variables and looping over steps if necessary.
